# Precartilage condensation during limb skeletogenesis occurs by tissue phase separation controlled by a bistable cell-state switch with suppressed oscillatory dynamics

**DOI:** 10.1101/2021.03.15.435301

**Authors:** T. Glimm, B. Kaźmierczak, C. Cui, S.A. Newman, R. Bhat

## Abstract

The tetrapod limb skeleton is initiated in unpatterned limb bud mesenchyme by the formation of precartilage condensations. Here, based on time-lapse videographic analysis of a forming condensation in a high-density culture of chicken limb bud mesenchyme, we observe a phase transition to a more fluidized state for cells within spatial compacted foci (protocondensations that will progress to condensations), as reflected in their spatial confinement, cell-substratum interaction and speed of motion. Previous work showed that galectin-8 and galectin-1A, two proteins of the galactoside-binding galectin family, are the earliest determinants of this process in the chicken limb bud, and that their interactions in forming skeletogenic patterns of condensations can be interpreted mathematically through a reaction-diffusion-adhesion framework. Based on this framework, we use an ordinary differential equation-based approach to analyze the core switching modality of the galectin reaction network and characterize the states of the network independent of the diffusive and adhesive arms of the patterning mechanism. We identify two steady states where the concentrations of both galectins are respectively, negligible, and very high. An explicit Lyapunov function shows that there are no periodic solutions. For sigmoidal galectin production terms, the model exhibits a bistable switch that arises from a monostable state via saddle-node bifurcation. Our model therefore predicts that the galectin network exists in low and high expression states separated in space or time without any intermediate states. This provides a causal basis for the observed outside vs. inside transition observed in the in vitro video data. We performed a quantitative analysis of the distribution of galectin-1A in cultures of condensing chick limb mesenchymal cells and found that the interior of the protocondensations had concentrations of this protein (compared to the immediate exterior) over and above that expected from its higher cell density, consistent with the model’s predictions. The galectin-based patterning network is thus suggested, on theoretical grounds, to incorporate a core switch independent of any spatial or temporal dynamics, that drives the chondrogenic cell state transition in limb skeletogenesis.

## 1 Introduction

Cell differentiation during embryogenesis has long been recognized to have features in common with multistable dynamical systems. This resemblance has provided motivation for mathematical and computational models of development over the past half-century (Kauffman (1969); Thom (1976); Keller (1994); Furusawa and Kaneko (2012); Huang (2012)). The analogy is appealing for several reasons: an organism’s genes (collectively its genome) and the RNA and protein molecules they specify can be conceived as a highly connected auto- and cross-regulatory multi-component system. Although the number of constituents (including alternative splice forms and covalently modified proteins) range in thousands in animals, for example, the number of distinct cell types are in the tens to hundreds (more than 250 in humans), suggesting that cell types may represent the discrete stationary states (time-independent solutions) or attractors of dynamical systems of gene regulatory networks (GRNs). Further, cell types are not only well-separated in the multidimensional state space defined by concentrations of gene products or transcription factors, they are also robust under perturbation, i.e., they exhibit stability in the face of physiological noise and transition between states along well-defined pathways during ontogeny or under experimental manipulation (Steinacher et al. (2016); Goode et al. (2016); Srivastava (2006)). All of this strengthens the analogy between cell types and the attractors of complex systems.

The analogy of developmental cell type transitions to the structure of dynamical systems also has severe limitations, however. The compositional differences (i.e., location in state space) between attractor states of a multicomponent dynamical system are a purely mathematical effect. Cell types, in contrast, embody coherent functions within their respective tissues: contractile muscles, supportive bones, excitable nerves and so forth. It is implausible that the suites of gene products required to support the coordinated tasks of any one cell type, let alone all of them, would arise simply as a consequence of dynamical partitioning (Newman (2020)). Moreover, the different cell types within an organ (e.g., in the lung, the gas permeable type 1- and surfactant-producing type-2 pneumocytes of the alveolar walls, and the capillary endothelium of the air sacs) need to operate as a unit (Ross et al. (2002)). It is highly unlikely that mathematically determined attractor states of a dynamical system would function in such a coordinated fashion (Newman (2020)).

Notwithstanding these caveats about the suitability of dynamical systems models of differences among cell *types*, in certain cases such methods have proved an apt modelling approach for the narrower concept of cell *state* transitions. Specifically, such models can capture important aspects of transitions between cell states adjacent to one another in a developmental lineage. Here, the states on either side of the transition have defined functional relationships that often depend on different levels of expression of a small number of key factors. Thus, dynamical systems, in the form of Boolean functions or ordinary differential equations (ODEs), have been used to model differentiation in the mammalian red and white blood cell-forming systems, (Chang et al. (2006); Mojtahedi et al. (2016)), regionalization of the gut in the sea urchin larva (Cui et al. (2017)), developmental cell type diversification in insect sensory organs and the nematode vulva (Corson et al. (2017); Corson and Siggia (2017)), the morphogenesis of flowers (Alvarez-Buylla et al. (2008)) and pluripotent stem cells exposed to various signaling factors (Saez et al., 2021).

Here, based on tracking cells in a living culture of precartilage cells undergoing the earliest event in skeletogenesis (what has been termed “compaction” (Barna and Niswander (2007)) or “protocondensation” (Bhat et al. (2011))), we find that this process involves a set of abrupt morphological and behavioral changes of the initially homogeneous cells. Specifically, the cells within the forming condensation became mainly confined within a sharp perimeter, less adherent to their substratum, and more rapidly moving than those immediately outside the focus. We wondered whether this was an example of a dynamical bifurcation in cell state, in this case leading to the incipient differentiation of the cartilaginous template of the avian limb skeleton.

In an earlier paper, we analysed the conditions for the breaking of spatial symmetry in an integro-differential equation model of the two-galectin reaction-diffusion-adhesion network of the developing avian limb (Glimm et al. (2014)). This “two galectins + ligands” (2GL) network comprised interactions of cells with the proteins galectin-1A (Gal-1A) and galectin-8 (Gal-8) and their galactoside-bearing cell surface counterreceptors or ligands. which experimentally had been found to mediate both protocondensation morphogenesis and the patterning of the subsequent precartilage condensations (Bhat et al. (2011)). Motivated by the transition in cell behavior observed in the cell tracking experiments, we focused here on the intracellular branch of this network, factoring out the cell-cell interaction branch required for spatial pattern formation. This has permitted us to study the dynamics of the transition between the intra-condensation and extra-condensation states in a reduced ODE system. Experimental evidence indicates that these states represent distinct cellular phenotypes at the biochemical, morphological, and functional levels, and not simply variant packing states or arrangements of a uniform cell type (Barna and Niswander (2007)). Our mathematical analysis supports this inference, indicating that the two-galectin system exhibits two stable states characterized by high and very low concentrations of the galectins.

Our model is thus an example of a dynamical system with switch-like behavior, arising (in an extended model with sigmoidal production terms) from a monostable system via a saddle-node bifurcation. This conceptually simple mechanism has been well-studied as a prototypical way in which molecular switches arise in regulatory networks, both in large-scale models (e.g. Tyson et al. (2001, 2003)) and more abstract geometric models motivated by Waddington’s epigenetic landscape analogy (e.g. Corson and Siggia (2017); Ferrell (2012); JutrasDubé et al. (2020), Saez et al. (2021)).

Although our model is based on experimentally determined interactions, it is relatively simple, arising from a reduction of a seven-species system of equations to a three-species system via elimination of “fast” variables (Glimm et al., 2014). As a consequence, it is analytically tractable – we are able to provide an explicit Lyapunov function, for instance. This has permitted us to infer that the core switch in the galectin-based skeletal patterning mechanism (which appears to have deep roots in the non-tetrapod gnathostomes (Bhat et al., 2016)), has no parametrically accessible oscillatory modes, despite the frequent association of these dynamical modes in biochemical and genetic networks (Goldbeter, 2018). This, along with the seemingly paradoxical foundation of solid skeletal tissues on tissue primordia that are fluidized during their initial differentiation (Jain et al., 2020; Kim et al., 2021) has implications for the evolution of limb skeletogenesis that are discussed in the concluding section.

## 2 Quantitative and qualitative differences in cell motion between intra-condensation and extra-condensation cells

In high-density (“micromass”) cultures of limb bud mesenchymal cells, initially homogeneous monolayers undergo focal rearrangements to form tightly packed condensations (Frenz and Newman, 1989). Before any overt cell rearrangement occurs, the protocondensations are marked, in the chicken system, by the expression of BMP receptor 1b and Gal-1A and Gal-8. We investigated the motility dynamics of single mesenchymal cells as protocondensations formed in vitro using cell-resolved, video-tracking techniques (Figure 1).

**Figure 1:**
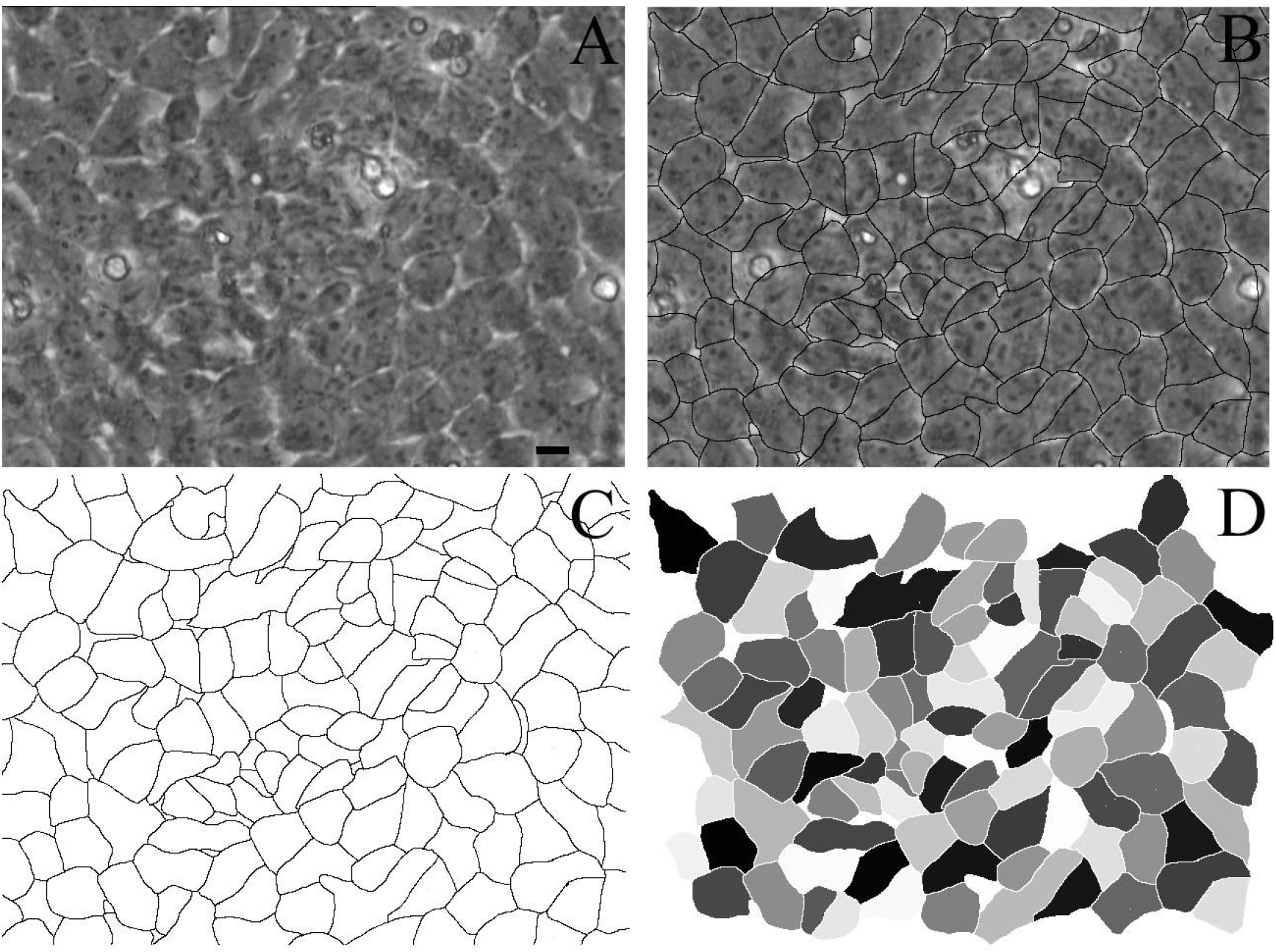
Image digitization. A, Original image: a single frame from the movie. B, Image with boundaries drawn by hand in Photoshop. C, Boundaries of tracked cells. D, Digitized image with gray-level-labeled cells. The scale bar in A represents a length of 10 *µ*m.

To record the 480 minutes during which the protocondensations emerged, we processed 25 successive images acquired at 20 min intervals. (We took time-lapse images at intervals of 2 minutes at a resolution of 640×480 pixels and chose every tenth image for processing. See movie file M1 in Supplemental Materials.) During the observation period, cells in the monolayer moved, changed their relative positions, and deformed continuously. In each image, we traced centers of condensation by determining the cell-substratum attachment area for each cell, smoothing this function over the whole image and then determining the contour that corresponded to a cell-substratum area of 160 *µ*m^2^ (Figure 2). Cells inside the condensation had smaller average contact area with the substratum than peripheral cells.

**Figure 2:**
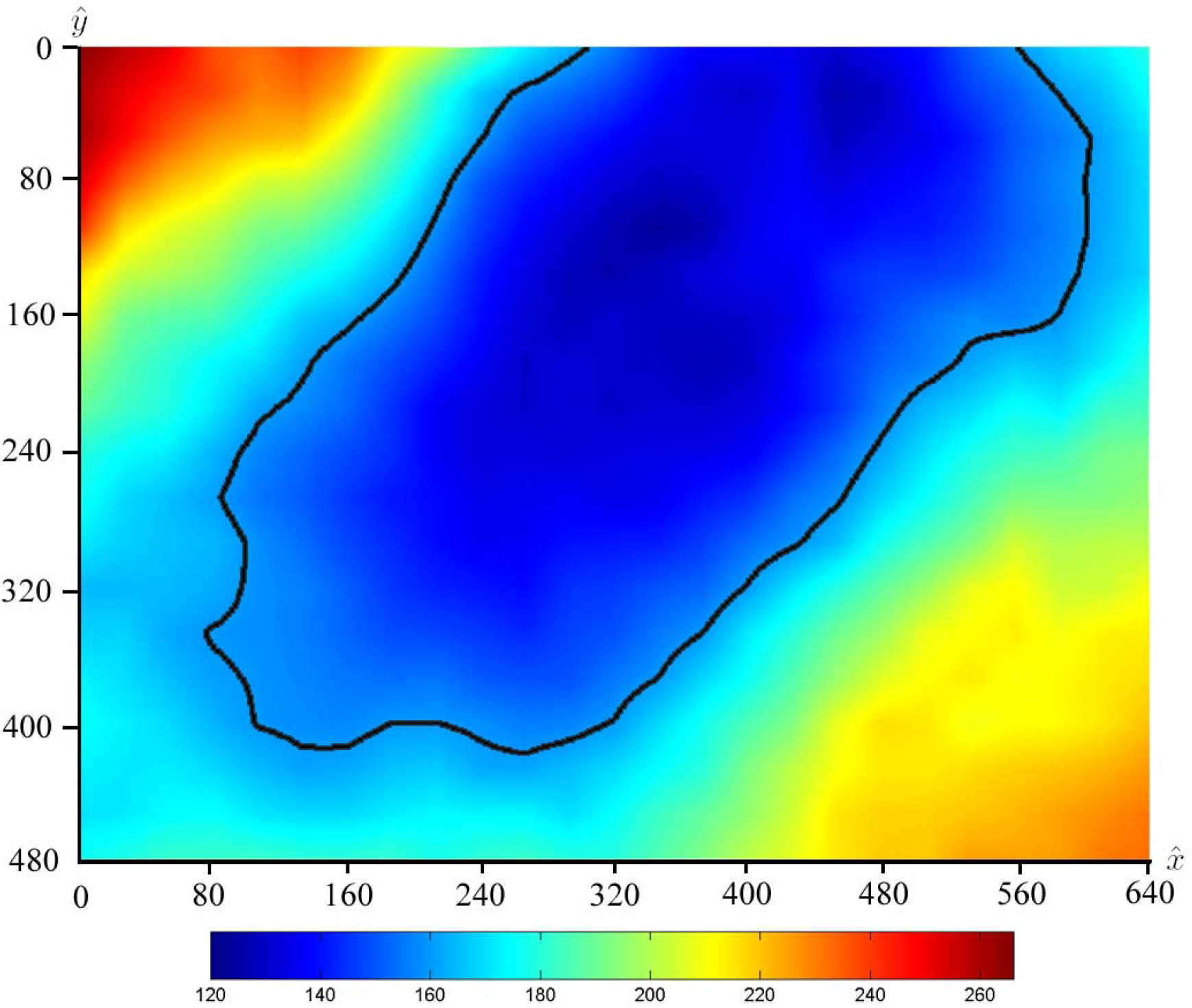
Heat map of average cell-substratum contact area for cells. See text for details. We characterized the condensation as the region bounded by contour line at average cell area = 160 *µ*m^2^ (black line). The units in the color bar are *µ*m^2^.

Of the 92 cells we tracked in the image set, 46 stayed exclusively inside the condensation center (i.e., within 57 *µ*m of the center), whereas 33 cells remained exclusively outside. The centers of mass of 13 cells crossed this boundary at least once in either direction during the 8 h of filming. The velocity statistics of the cells residing exclusively inside and outside the forming condensation are shown in Table 1.

**Table 1:** Comparison of velocities of cells residing exclusively inside and outside precartilage condensation.

|  | Inside | Outside |
| --- | --- | --- |
| Number of cells | 46 | 33 |
| mean value ($\mu\text{m}/\text{min}$) | 0.1141 | 0.1012 |
| standard deviation | 0.0761 | 0.062 |
| median ($\mu\text{m}/\text{min}$) | 0.0941 | 0.0867 |

The mean speed of cells exclusively inside condensation foci exceeded that of cells that remained exclusively outside, and the same held true for the median value. (A two-tailed *z*-test to test the significance of the difference of the means yielded a *p*-value of approximately 10^−4^.) A total of 615 out of the 1104 measured speeds for cells staying exclusively inside the condensation center (or 55.71%) were greater than the median value (0.0867 *µ*m*/*min) of the speeds of cells remaining outside the condensation center.

Consistent with this difference in mean speeds the mean squared displacement (MSD) of cells inside condensation centers increased with time at a higher rate than that of cells outside condensations (see Figure 3). Since the slope of the MSD is proportional to the diffusion coefficient (diffusion understood as randomly directed cell movement), this means that cells within condensations diffused faster than those outside.

**Figure 3:**
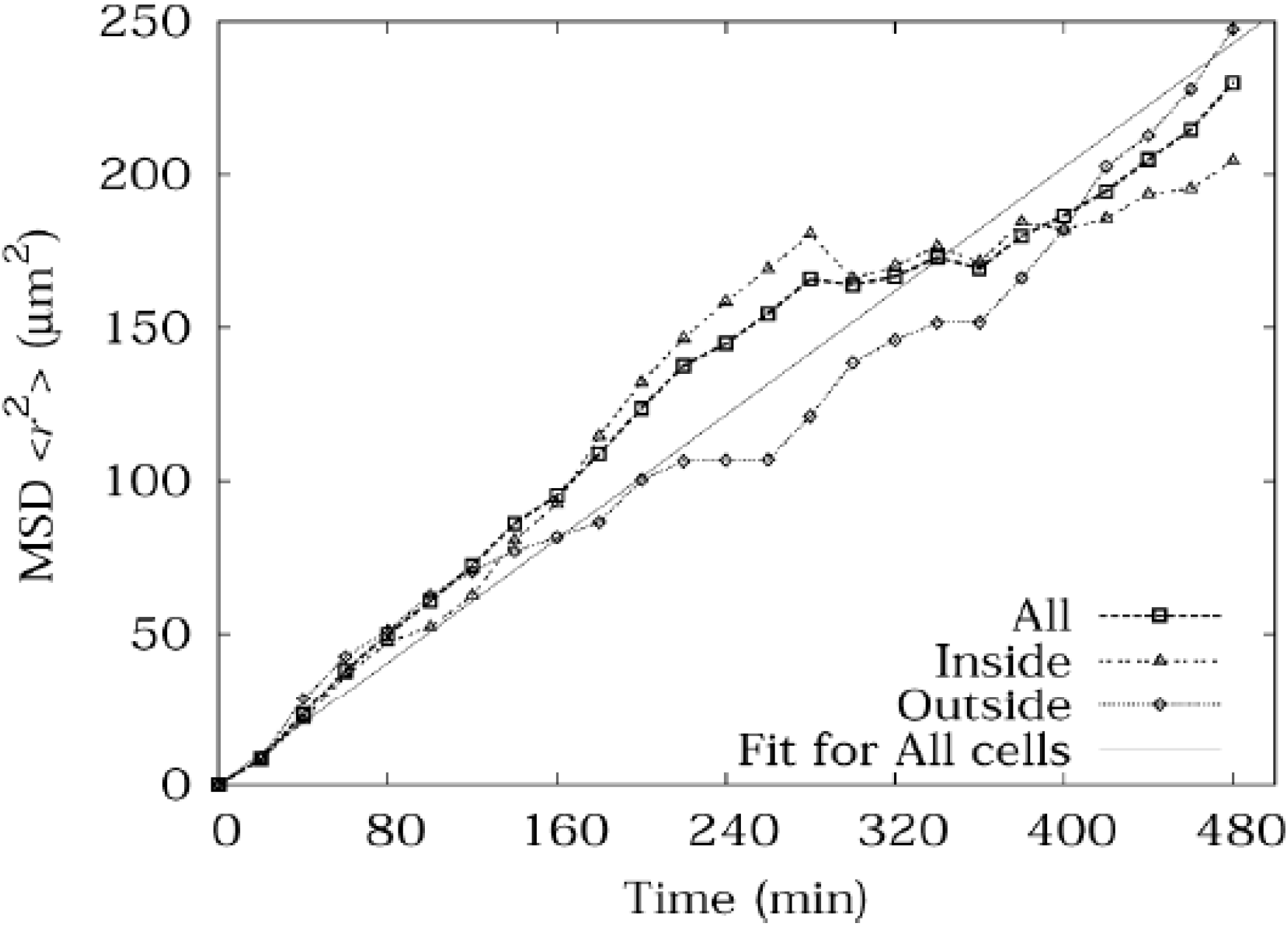
Mean-squared displacement of all cells, cells inside the condensation center all the time and cells outside the condensation center all the time, with center-of-mass movement of all cells and drift of cells outside the condensation center removed. The overall diffusion coefficient of cells was determined as 0.5058±0.011*µ*m^2^*/*min.

We characterized the linear dimension and number of cells in the observed condensation and found that interior cells in this region had smaller areas of contact with their substratum than peripheral cells. While cells inside and far outside the forming condensation have random speeds and directions of motion, cells in the periphery of a condensation tend to move towards the center of the condensation (see Figure S1 in the Supplementary Material). This is consistent with a hypothetical haptotaxis mechanism, with no intrinsic directionality, but based on random cell movement with preferential sticking to increasingly adhesive substrata. Such a model does not depend on, or imply, a qualitative difference between intra-condensation and extra-condensation mesenchymal cells (Dickinson 2000).

However, contrary to the predictions of the haptotaxis model, we found (as described above) that cells within the forming condensation moved significantly faster than those in the periphery. The altered cell-substratum adherence and motile behavior of the cells in the developing condensation relative to those in the periphery, and the general confinement of those cell within a well-defined boundary, suggests that the mesenchymal tissue inside the focus undergoes a transition to a more fluid state (Jain, 2000; Kim, 2021).

## 3 The spatially independent core network of the 2GL model

### 3.1 Derivation

To determine whether the dynamics of the 2GL network can provide insight into the observed phase transition and separation, we considered a version of the model in which the assumption of fast galectin-counterreceptor binding reduces the seven variables of the network and their interactions to a system of four coupled ODEs.

The galectin dynamical network, based on experimental data described in Bhat et al. (2011), is represented schematically in the diagram in Figure 4. The associated spatiotemporal partial differential equation model of Glimm et al. (2014) also includes morphogen diffusion and cell adhesion and motion, but not a detailed analysis of the galectin network by itself. The main constituents and regulatory interactions of the model of Glimm et al. (2014) are summarized as follows: There are two freely diffusible, ECM-bound proteins, Gal-1A and Gal-8. They bind to two different cell-membrane-bound counterreceptors. The first of these (“Gal-8 counterreceptor”) can only bind to Gal-8, which activates the production of Gal-1A. The second (“shared counterreceptor”) can bind to both Gal-1A and Gal-8. The binding of Gal-1A to the shared counterreceptor activates both the production of Gal-8 and the shared counterreceptor itself. The binding of Gal-8 to the shared counterreceptor has no biochemical regulatory effect, but interfere with the binding, and hence action, of Gal-1A.

**Figure 4:**
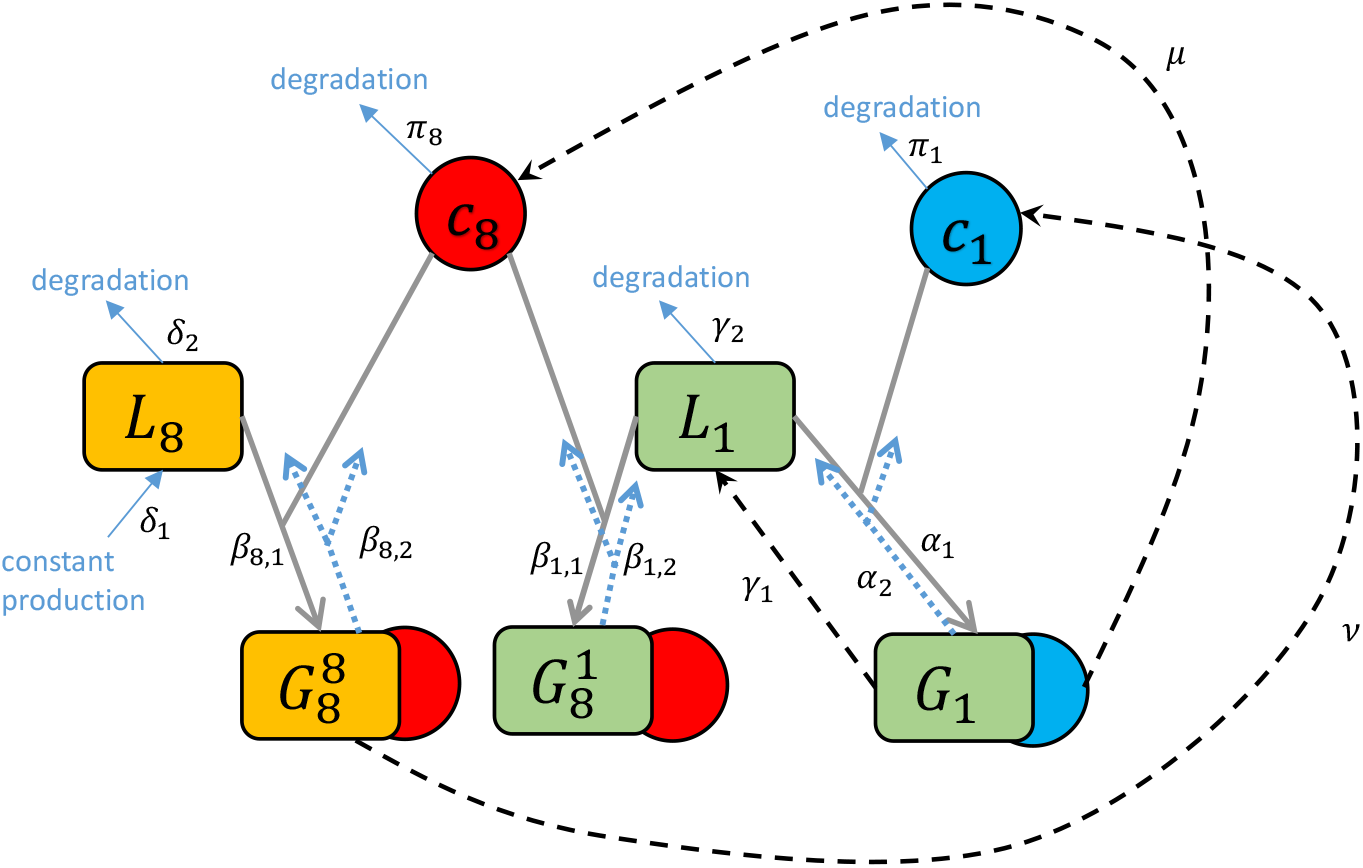
Schematics of the galectin network. Galectins are denoted by *c*_1_ and *c*_8_, unbound counterreceptors by *L*_8_ and *L*_1_ and the complexes of galectins bound to counterreceptors by 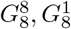 and *G*_1_. Solid arrows denote reversible binding of galctins and counterreceptors with the unbinding shown by dotted arrows. Dashed arrows denote activation.

We denote the concentration of unbound shared counterreceptors by *L*_1_ and *L*_8_ is the concentration of Gal-8’s counterreceptor. *G*_1_ is the concentration of the complex of Gal-1A and the shared counterreceptor, 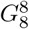 is the concentration of the complex of Gal-8 and its counterreceptor, and 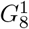 is the concentration of the complex of the shared counterreceptor with Gal-8. The concentrations of the two unbound galectins are denoted by *c*_1_ and *c*_8_, respectively. Note that these are secreted proteins, so in principle they can diffuse through the extracellular matrix. Thus in spatially extended systems, galectins provide signals that can coordinate cell behavior over distances. However in the present paper, we are interested only in the local properties of the galectin regulation network and its possible states.

Based on experimental data in Bhat et al. (2011) and using simple mass action laws in a compartmental model (Figure 4), we can set up the following system of ODEs to describe the dynamics. (These equations were derived in Glimm et al. (2014) from a spatiotemporal model under the assumption of zero cell motility.)

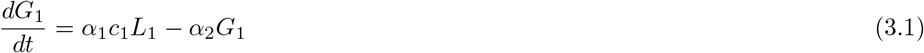

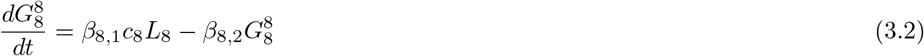

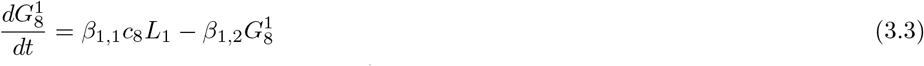

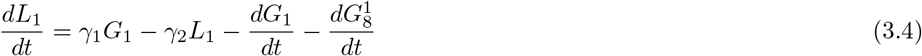

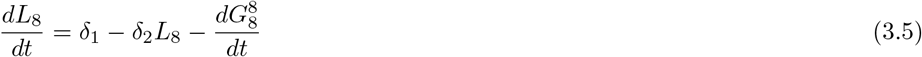

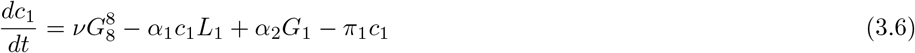

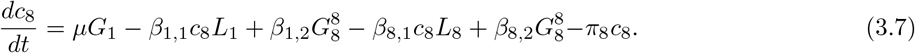

We make the assumption that binding of galectins occurs on a faster time scale than production of counter-receptors and galectins. This is expressed by the quasi-steady state assumptions

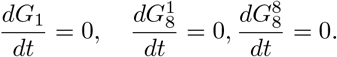

This makes *G*_1_,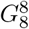 and 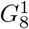 effectively functions of the remaining variables, namely

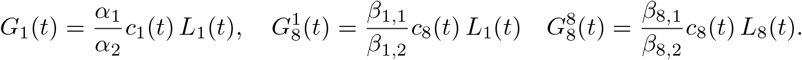

The remaining variables then satisfy

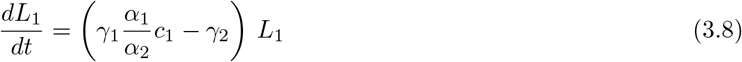

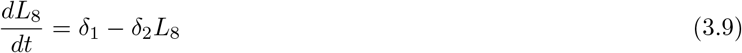

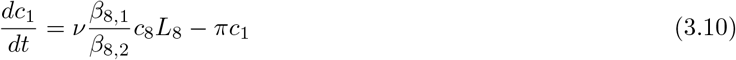

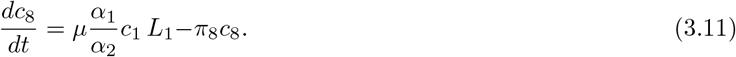

We introduce nondimensional variables 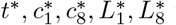 via

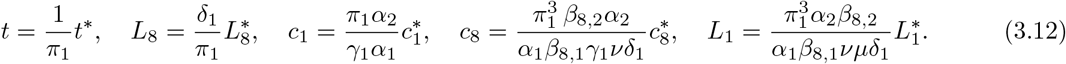

We define dimensionless parameters

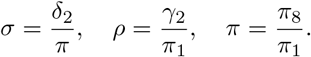

Rewriting the system (3.8)–(3.11) with these new dimensionless variables and dropping the stars, the system (3.8)–(3.11) is

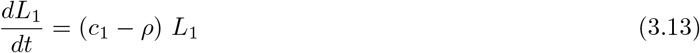

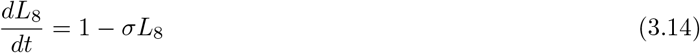

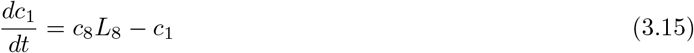

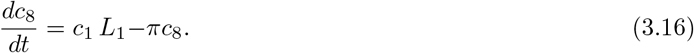

This is the central system of this paper. Note that the equation for *L*_8_(*t*) is independent of the other variables and can be solved explicitly as *L*_8_(*t*) = 1*/σ* + (*L*_8_(0) − 1*/σ*) exp(−*σ t*). In particular, *L*_8_(*t*) → 1*/σ* as *t* → ∞. This is why we will also sometimes consider the three variable system where *L*_8_ = 1*/σ*:

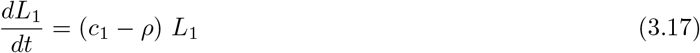

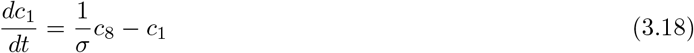

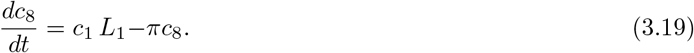

### 3.2 Parameters

The system of equations (3.13)–(3.16) contains three parameters *ρ, σ* and *π*. All three represent the ratios of degradation constants which can in principle be estimated based on experimental data for the degradation rates of galectins and membrane-bound receptors. (See Table 2; see also the list of parameters used by Bhat et al. (2019).) A crucial role is played by the time scale 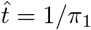 of the nondimensionalized time variable *t*^∗^ in (3.12). The half life of galectins is unknown, but likely on the order of several hours (Chen et al. (2014)). Based on the assumption of *π*_1_ ≈ 1*/*day (Bhat et al. (2019)), which corresponds to a half life of Gal-1A of about 16 hours, this gives as the time scale a value of 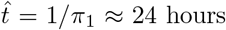. Likewise, the half-life of counterreceptors has not been measured directly, but an analysis of 552 human cell-surface glycoproteins yielded a mean half-life of 19.6 hours, with glycoproteins with receptor functions tending to have smaller half-lives than others (Xiao and Wu (2017)). This means that the half-lives of galectins and their counterreceptors are likely of the same order of magnitude. Our mathematical results are independent of the ratio of half lives, but for the illustrations, we assume a ratio of 2.

**Table 2:** Parameters for the system of equations (3.13)–(3.16). Plausible ranges are within one order of magnitude of the estimated value. Note that due to the molecular similarity of Gal-1A and Gal-8 on the one hand, and of the counterreceptors on the other, it is plausible to assume that *π* and the ratio *ρ/σ* are both close to unity.

| parameter | definition | meaning | est. value | est. plausible range |
| --- | --- | --- | --- | --- |
| $\rho$ | $\gamma_2/\pi_1$ | ratio of half-lives of Gal-1A and shared counterreceptor | 2 | $0.2 < \rho < 20$ |
| $\sigma$ | $\delta_2/\pi_1$ | ratio of half-lives of Gal-1A and Gal-8's counterreceptor | 2 | $0.2 < \sigma < 20$ |
| $\pi$ | $\pi_8/\pi_1$ | ratio of half-lives of Gal-1A and Gal-8 | 1 | $0.1 < \pi < 10$ |

### 3.3 Positivity of solutions

The system (3.13)–(3.16) preserves positivity of solutions, consistent with the constraint that negative values for the variables are not biologically meaningful:

#### Theorem 3.1.

*Suppose* (*L*_1_(*t*), *L*_8_(*t*), *c*_1_(*t*), *c*_8_(*t*)) *is a solution to the system (3*.*13)–(3*.*16) with positive initial condition: L*_1_(0), *L*_8_(0), *c*_1_(0), *c*_8_(0) *>* 0. *Then L*_1_(*t*), *L*_8_(*t*), *c*_1_(*t*), *c*_8_(*t*) *>* 0 *for all t >* 0.

*Proof*. It is clear that *L*_8_(*t*) will be positive for all *t >* 0 if *L*_8_(0) is positive. Note that *L*_1_(*t*^∗^) = 0 for some time *t*^∗^ implies that *L*_1_(*t*) = 0 for all time *t*. Moreover, by inspection 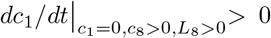 and 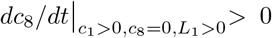, so that neither *c*_1_ nor *c*_8_ can become zero when all other variables are positive. The only remaining possibility is that both *c*_1_ and *c*_8_ become zero at the same time, but then note that *c*_1_(*t*^∗^) = *c*_8_(*t*^∗^) = 0 for some time *t*^∗^ implies that *c*_1_(*t*) = *c*_8_(*t*) = 0 for all times *t*.□

## 4 Phase space

### 4.1 Analysis

The system (3.13)–(3.16) has two steady states: besides the (mathematically, though not biologically - see below) trivial one 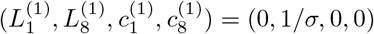, there is one nontrivial state

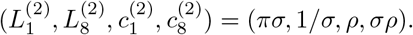

The trivial steady state is always stable; in fact, its linearization matrix has the four negative eigenvalues −*ρ*, −*σ*, −1 and −*π*. The linearization matrix at the nontrivial steady state 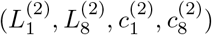 is given by

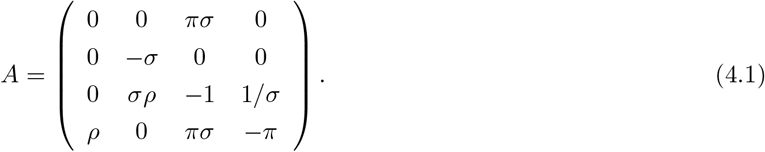

Its characteristic polynomial is given by

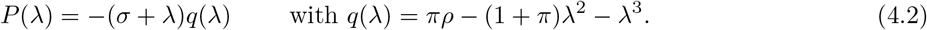

Thus, one of the eigenvalues is always *λ*_1_ = −*σ*. The other eigenvalues are the roots of *q*(*λ*). A straightforward application of the Routh-Hurwitz stability criterion confirms that *q*(*λ*) has exactly one root with positive real part. The other two roots must have negative real parts.

Thus the nontrivial steady state is always a saddle with the eigenvalue signature (+, −, −, −). We note that the equation (3.14) can be solved explicitly to give *L*_8_(*t*) = (1 − exp(−*σt*)*/σ* + *L*_8_(0) exp(−*σt*)). It is helpful to set *L*_8_ = 1*/σ* to obtain a three dimensional phase space that can be visualized. The phase space in Figure 5, typical for this system, is separated into two three-dimensional regions: One region where trajectories converge towards the trivial steady state 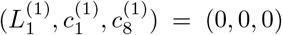 and one where trajectories diverge to infinity in all three components. The separatrix is the two-dimensional stable manifold of the saddle point 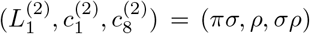. An analogous behavior in two dimensions is displayed in an illustrative simplified system with only two variables in Appendix B; see in particular Figure 10.

**Figure 5:**
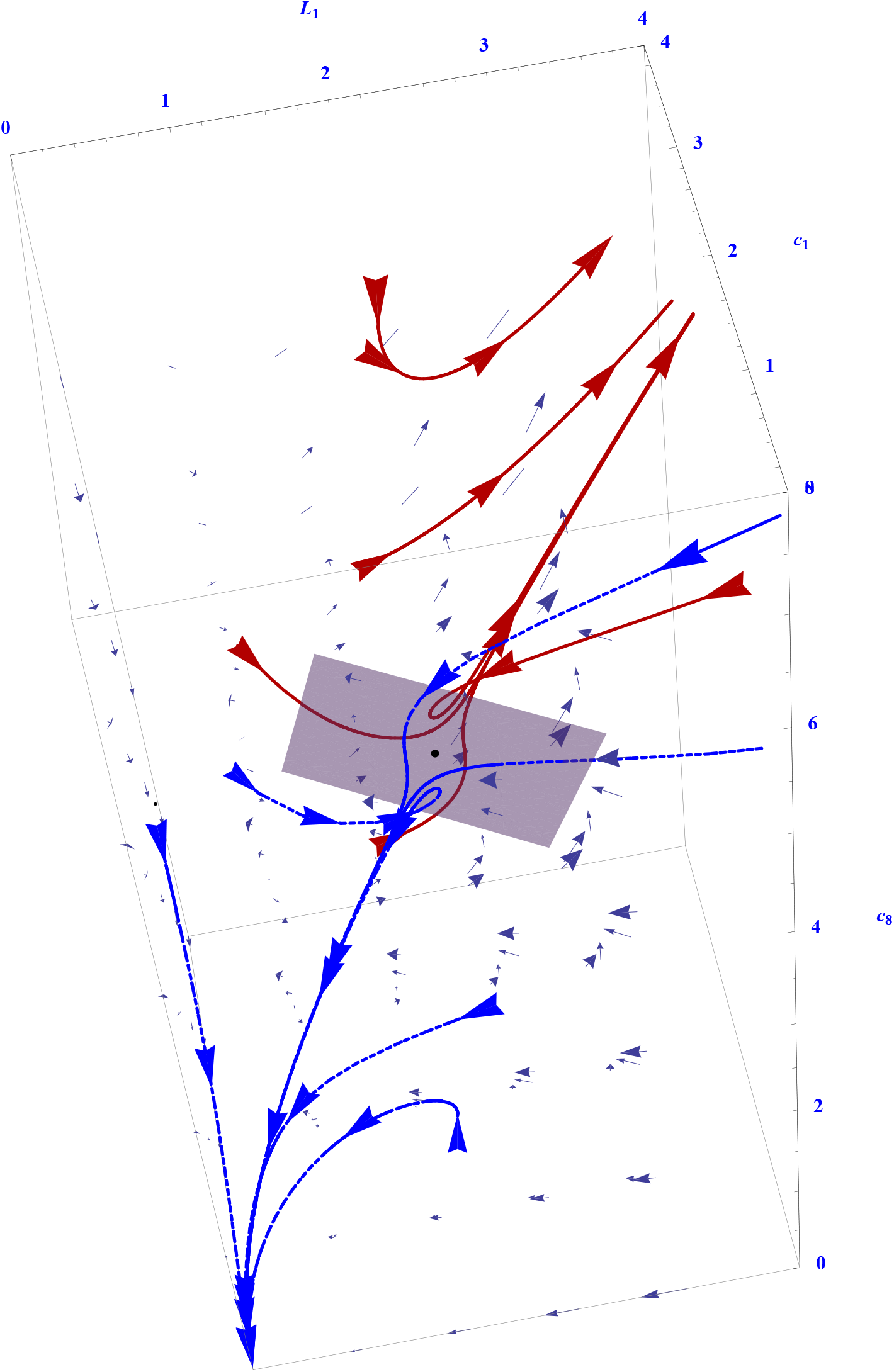
Phase space for the system of equations (3.13)–(3.16) for *σ* = *ρ* = 2 and *π* = 1. Blue dashed trajectories converge to the trivial steady state. Red solid trajectories diverge to infinity. The saddle is shown with tangent space to the stable manifold.

This simple analysis indicates that there are two stable states: the trivial steady state, where all concentrations are zero, as well as a second state, in which concentrations grow without bounds. (See Corollary 4.3 for a mathematical proof that solution curves are in fact either unbounded or converge to one of the two equilibria, the stable trivial one or the saddle.) Biologically, these correspond to the states of mesenchymal cells outside and within the protocondensations, respectively.

The first state also corresponds to the network state wherein the Gal-1A and Gal-8 mRNA levels are sparse. This was indeed observed in the cells that are peripheral to protocondensations by in situ hybridization (Bhat et al, 2011). Cells that were included in protocondensations showed high levels of Gal-1A and Gal-8 mRNA levels and therefore represented the second state.

This core network was mathematically abstracted from the full experimentally founded 2GL pattern-forming system. In the full biological context, additional layers of spatiotemporal regulation will prevent the concentrations of a network’s components from growing out of bounds. For example, the network has been shown to be under negative regulation by Notch signaling (Bhat et al. (2019)).

### 4.2 Absence of substantial oscillations

Oscillatory behavior is often found in gene regulatory networks and plays a crucial role, for instance, in Kaneko’s theory of cell differentiation (Suzuki et al. (2011)). Temporal oscillations of the expression rates of the transcription factor Hes1 occur during condensation formation in limb development, where they interact with the 2GL network (Bhat et al. (2019)). It is thus natural to ask whether the system (3.17)–(3.19) can exhibit such oscillatory behavior.

In this section, we prove that this is not the case. In fact, solutions are either unbounded, or they converge to either the stable trivial steady state or the saddle point. This follows from the existence of a Lyapunov function:

#### Theorem 4.1.

*Consider the function*

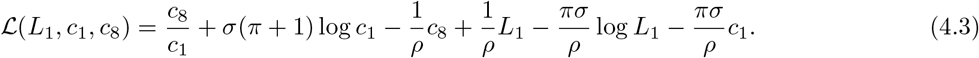

*Then* ℒ (*L*_1_, *c*_1_, *c*_8_) *is decreasing along solutions* (*L*_1_(*t*), *c*_1_(*t*), *c*_8_(*t*)) *of the system (3*.*17)–(3*.*19). Indeed, for* (*L*_1_(*t*), *c*_1_(*t*), *c*_8_(*t*)) *>* 0, *we have*

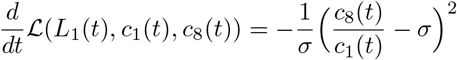

*Proof*. This follows directly from computing the derivative of ℒ and substituting the derivatives for *c*_1_, *c*_8_, *L*_1_ and *L*_8_ with the equations (3.17)–(3.19).□

*Corollary* 4.2. There are no non-constant periodic solutions to the system (3.17)–(3.19).

*Proof*. According to Theorem 3.1, we can assume that *c*_1_(*t*) ≠ 0 for all *t* ∈ [0, *T*). If (*L*_1_(·), *c*_1_(·), *c*_8_(·)) is a periodic solution of period *T >* 0, then

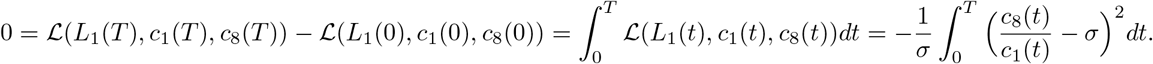

It follows that for such a solution we would have *c*_8_(*t*) = *σc*_1_(*t*) for all *t* ∈ [0, *T*]. From the second equation, this would imply that *c*_1_(*t*) = *c*_1_(0) and *c*_8_(*t*) = *c*_1_(0)*σ* for all *t* ∈ [0, *T*]. Then the left hand side of the third equation would be zero, implying that *L*_1_(*t*) = *const*. But then from the first equation we obtain *c*_1_ = *ρ*, hence *L*_1_ = *πσ*. Hence the solution coincides with the unstable stationary point. □

*Corollary* 4.3. If (*L*_1_(·), *c*_1_(·), *L*_1_(·)) is a bounded solution, then either (*L*_1_(·), *c*_1_(·), *L*_1_(·)) → (0, 0, 0), or (*L*_1_(·), *c*_1_(·), *L*_1_(·)) → (*πσ, ρ, σρ*) as *t* → ∞.

*Proof*. ℒ (*L*_1_(*t*), *c*_1_(*t*), *c*_8_(*t*)) is decreasing on trajectories, so

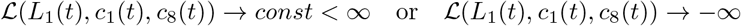

as *t* → ∞. In the first case *d*ℒ*/dt* → 0 as *t* → ∞, so 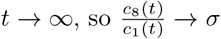. This means that (*L*_1_(*t*), *c*_1_(*t*), *c*_8_(*t*)) approaches the intersection of a level set of ℒ (*L*_1_, *c*_1_, *c*_8_) with the plane *c*_8_ = *σc*_1_. Since (*L*_1_(*t*), *c*_1_(*t*), *c*_8_(*t*)) is bounded and there are no periodic orbits according to Corollary 4.2, this implies that (*L*_1_(*t*), *c*_1_(*t*), *c*_8_(*t*)) must converge to one of the steady states, either (0, 0, 0) or the saddle point (*πσ, ρ, σρ*) In the second case *c*_1_(*t*) cannot be bounded away from zero. This means *L*_1_(*t*) is not bounded away from zero either. By inspection of the system (3.17)–(3.19), this implies that (*L*_1_(*t*), *c*_1_(*t*), *c*_8_(*t*)) will eventually lie in the basin of attraction of the nodal sink (0, 0, 0). Thus (*L*_1_(*t*), *c*_1_(*t*), *c*_8_(*t*)) → (0, 0, 0) as *t* → ∞.□

While there are no limit cycles, there can be damped spirals in the vicinity of the saddle point. This is the case for complex eigenvalues:

#### Proposition 4.4.

*The saddle point of system (3*.*13)–(3*.*16) has a pair of complex eigenvalues if and only if the parameters ρ and π satisfy*

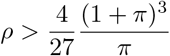

*Proof*. Consider the characteristic polynomials (4.2) of the linearization matrix (4.1). The discriminant of the third order polynomial *q*(*λ*) is given by Δ = 4*π*(1 + *π*)^3^*ρ* − 27*π*^2^*ρ*^2^. Complex eigenvalues exist if and only if Δ *<* 0. This is equivalent to the above inequality. □

If the saddle point is a spiral saddle with complex eigenvalue *λ*, then the ratio *b* = |Im(*λ*)*/*Re(*λ*)| is a measure of how tightly the spiral is coiled: Close to the saddle point, the projection of solution curves onto the stable manifold is a spiral, whose distance to the saddle decreases by the factor exp(−2*π/b*) with each turn. Thus in the limit *b* → ∞, the spiral approaches an ellipse, and in the limit *b* → 0, it approaches a straight line. A graph of *b* = |Im(*λ*)*/*Re(*λ*)| as a function of the parameters *ρ* and *π* shows that *b* is an increasing function of *π* for fixed *ρ* (see Figure 6). However, it is bounded from above:

**Figure 6:**
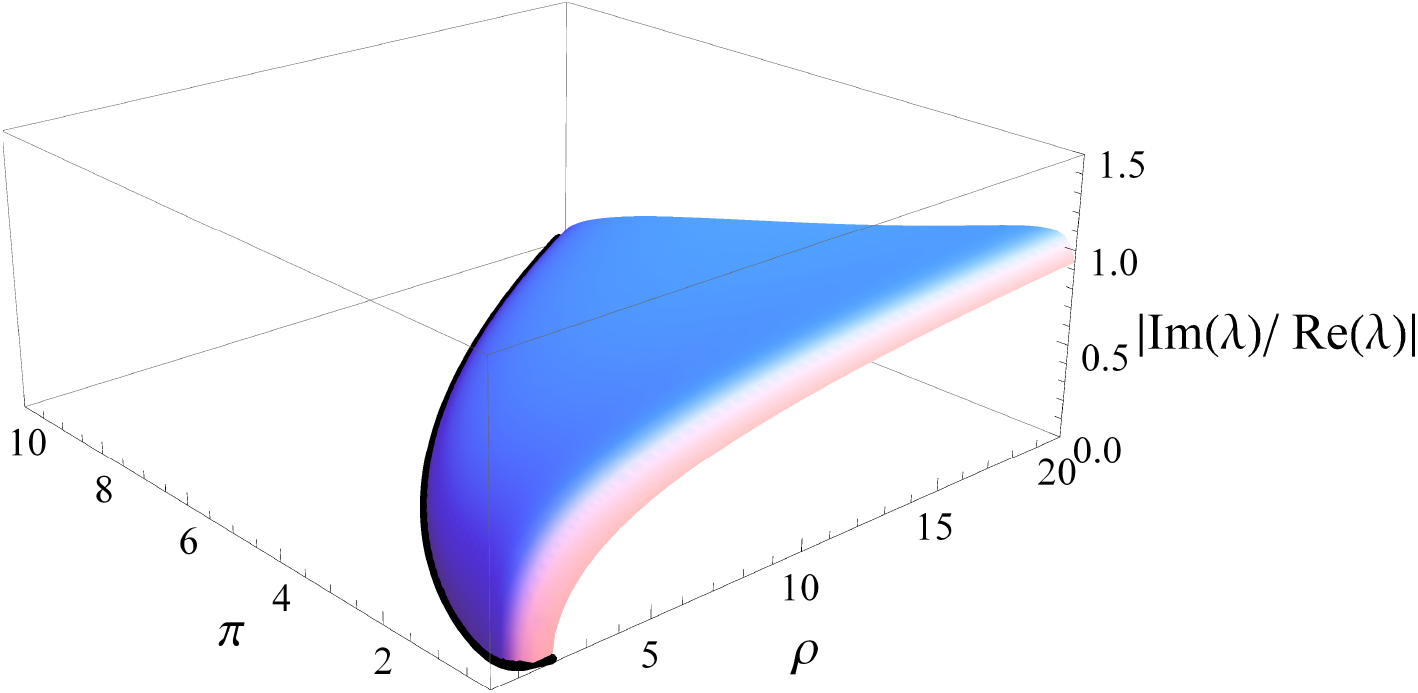
Plot of the ratio —Im(*λ*)*/*Re(*λ*)— of the complex eigenvalue of the saddle point of (3.13)–(3.16) as a function of the parameters *ρ* and *π*.

#### Proposition 4.5.

*Let λ*(*ρ, π*) *and* 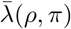 *denote the pair of complex eigenvalues of the saddle point of the system (3*.*13)–(3*.*16) which exist for* 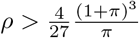 *Then for fixed π, we have*

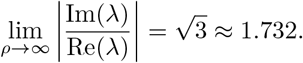

*Proof*. See Appendix C. □

The result shows that there is a lower limit on the tightness of the spiral, implying that from a practical point of view, oscillatory behavior does not play a role for any parameter values of the system (3.13)–(3.16).

### 4.3 An illustrative modification: Sigmoidal production terms

Note that the system (3.13)–(3.16) does not exhibit bifurcations under variation of the parameters. However, its phase portrait has two stable states separated by a separatrix surface, the stable manifold of a saddle point. This is analogous to the phase portrait of a bistable system (i.e., a switch) that arises from a monostable system via a saddle-node bifurcation. An analogous role to the second stable node in such a system is played by trajectories which go to infinity in our model, so that we have two possible long term states with attracting basins of nonzero measure: the zero steady state and a second one represented by trajectories going to infinity.

As a subsystem of the full developmental mechanism, the model’s variables are isolated from regulatory interactions that would curtail such extreme behaviors. We have therefore investigated a version of the model with some additional biologically motivated assumptions that can plausibly stand in for the complexity of the natural setting: Instead of basing the production rates of galectins and counterreceptors on mass action kinetics, we substitute Michaelis-Menten kinetics. This means that production rates do not simply increase proportionally to counterreceptor concentrations without bounds, but that they approach a finite maximum. With this modification the concentrations of the galectins cannot increase without bounds, but reach a second steady state. This yields the following model:

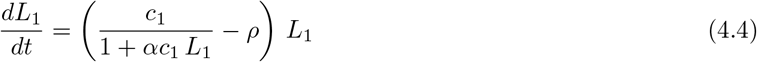

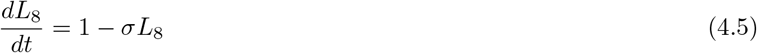

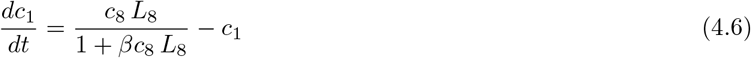

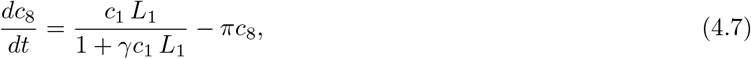

where *α, β* and *γ* are additional (nonnegative) parameters. Note that the model (3.13)–(3.16) corresponds to *α* = *β* = *γ* = 0. For fixed *α, β* and *γ*, there are pairs of solutions of the steady state equations given by

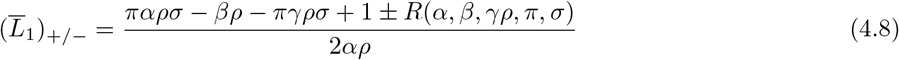

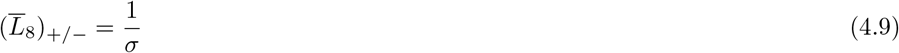

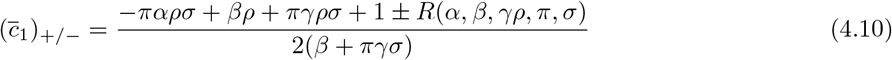

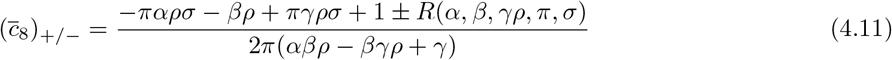

where

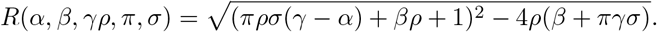

Here the subscript “+” denote the larger branch and “−” the smaller branch. In contrast to the simpler “mass action” system (3.13)–(3.16), this system does undergo a saddle-node bifurcation when *R*(*α, β, γρ, π, σ*) switches from imaginary to real values such that both branches of the steady states (4.8)–(4.11) are positive. The larger branch 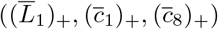 is stable, the lower branch 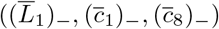 is a saddle. An example of a bifurcation diagram is shown in Figure 7.

**Figure 7:**
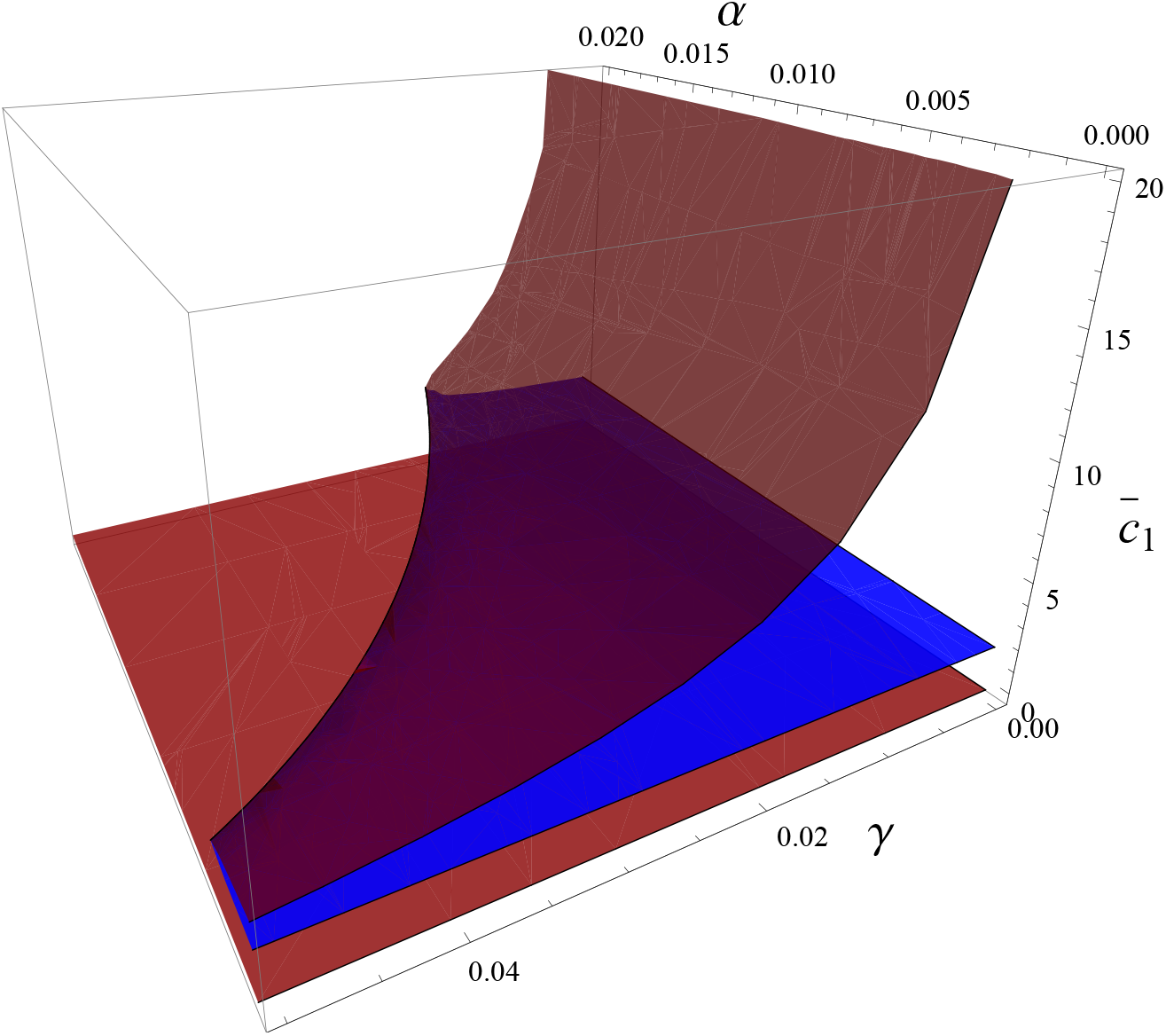
Bifurcation diagram for the equations (4.4)–(4.7) for *π* = 1, *σ* = *ρ* = 2 and *β* = 0.01. The vertical axis shows the steady state 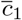. Stable nodes are shown in red. Saddles are shown in blue. Note the saddle-node bifurcations along a curve in *α*–*γ* space.

**Figure 8:**
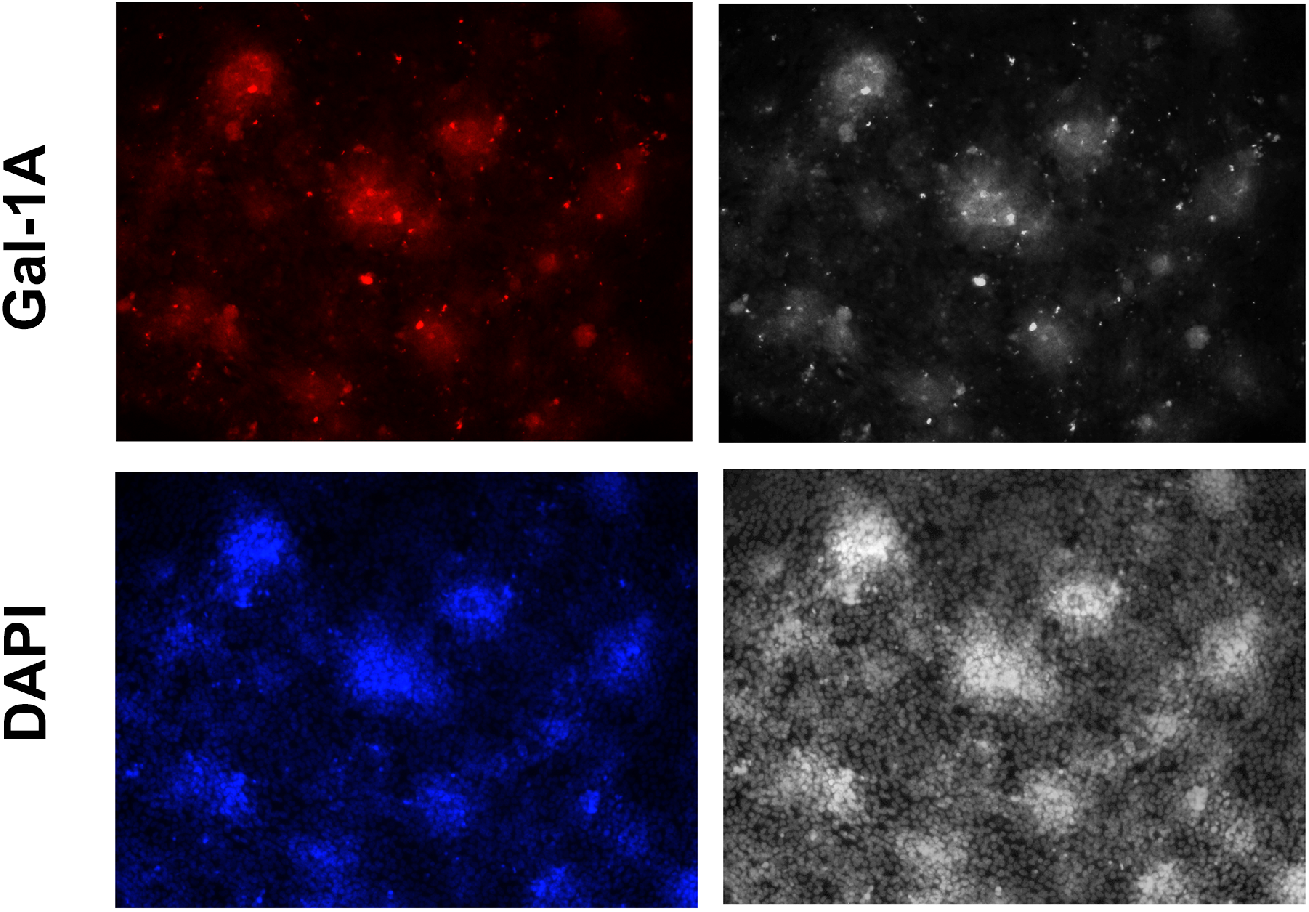
High resolution immunofluorescence images of precartilage condensations imaged for Gal-1A (top row) and DNA (using DAPI (bottom row)). Left column: Original images. Right column: grayscale intensity counterparts of the left column images.

Note that the simpler “mass action” system (3.13)–(3.16) can be formally obtained from the system (4.4)– (4.7) via setting *α* = *β* = *γ* = *ε* and taking the limit *ε* → 0. Indeed, one checks that with *α* = *β* = *γ* = *ε*, one gets

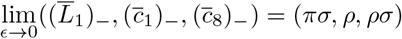

and

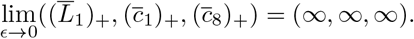

One thus recovers the “infinite” state and the saddle of the system (3.13)–(3.16) as the limits of the stable node and the saddle point of the system (4.4)–(4.7), respectively.

We emphasize that the sigmoidal production terms in this modification of the core model are not themselves based on experimental results. They serve as a heuristic substitute for the ancillary regulatory networks in which the core cell-state switch of the 2GL mechanism (itself experimentally based) is embedded in its natural developmental setting. Such interactions. essential to the switch’s functioning in spatiotemporal pattern formation would, similarly to the sigmoidal terms, prevent the attainment of of the extreme states its specifies on its own.

## 5 Testing the model

In contrast to a spectrum of biochemical states with progressively changing galectin expression, our model predicts the concurrent emergence of two distinct dynamical cell states: one with ‘high’ galectin production (‘intra-protocondensation cells’) and the other with ‘low’ galectin production rates (‘extra-protocondensation cells’) as a consequence of the reaction kinetics of the galectin network.

Although we had previously observed an increase in the concentration of both galectins accompanying precartilage protocondensation in vitro (Bhat et al., 2011), the bistable behavior exhibited by the spatially independent core network motivated a more quantitative evaluation, We therefore acquired high resolution images of condensing mesenchyme in high density chick limb bud micromass cultures. Using fluorescent immunostaining for Gal-1A (with counterstaining for DNA using DAPI), we found that cells within forming protocondensations had significantly higher levels of Gal-1A compared to cells outside. However, these differences are accompanied by an increase in cell density as condensation proceeds, and for the present purposes it was important to separate the effects of higher Gal-1A expression from an artifactual overestimation of its concentration due to spatially restricted cellular compaction.

To address this, we plotted the mean intensity of Gal-1A signal against the intensity of the DAPI signal, yielding a diagram that approximates Gal-1A concentration as a function of cell density (Figure 9). If all cells expressed Gal-1A at the same rate, we would expect Gal-1A concentration to be a linear function of cell density. Instead, the resulting graph was convex and a least squares approximation yielded an almost exactly quadratic curve (exponent 2.03). Therefore, the levels of Gal-1A concentration *normalized by cell density* itself increased approximately linearly with cell density, indicating significantly higher per cell expression within protocondensations than outside them.

**Figure 9:**
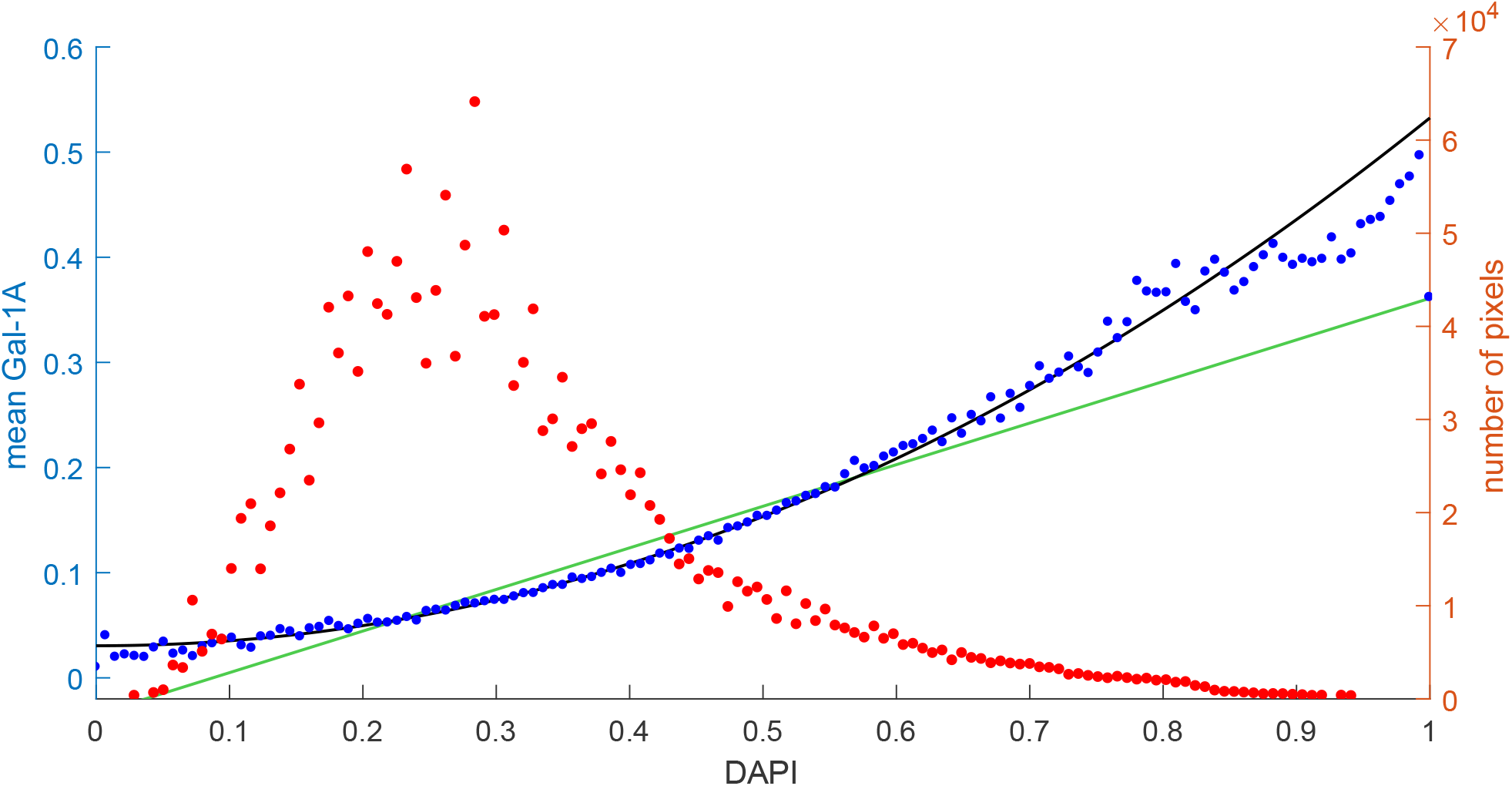
Intensity of the grayscale Gal-1A image as a function of the intensity of DAPI. Values are scaled from 0 (completely black pixels) to 1 (white pixels of maximum intensity). Blue dots indicate the mean Gal-1A intensity of pixels with the corresponding DAPI intensity. Red dots indicate the number of pixels for each DAPI intensity, showing that most pixels have an intensity between about 0.1 and 0.5. Green straight lline: linear regression line of the Gal-1A data, weighted by number of pixels, given by Gal-1A = −0.034623 + 0.3956 DAPI. Black curve: Nonlinear least squares regression of the Gal-1A data of the form Gal-1A = *a* + *b* DAPI^*c*^ weighted by number of pixels. Best-fit parameters are *a* = 0.030523, *b* = 0.50199, *c* = 2.0298.

**Figure 10:**
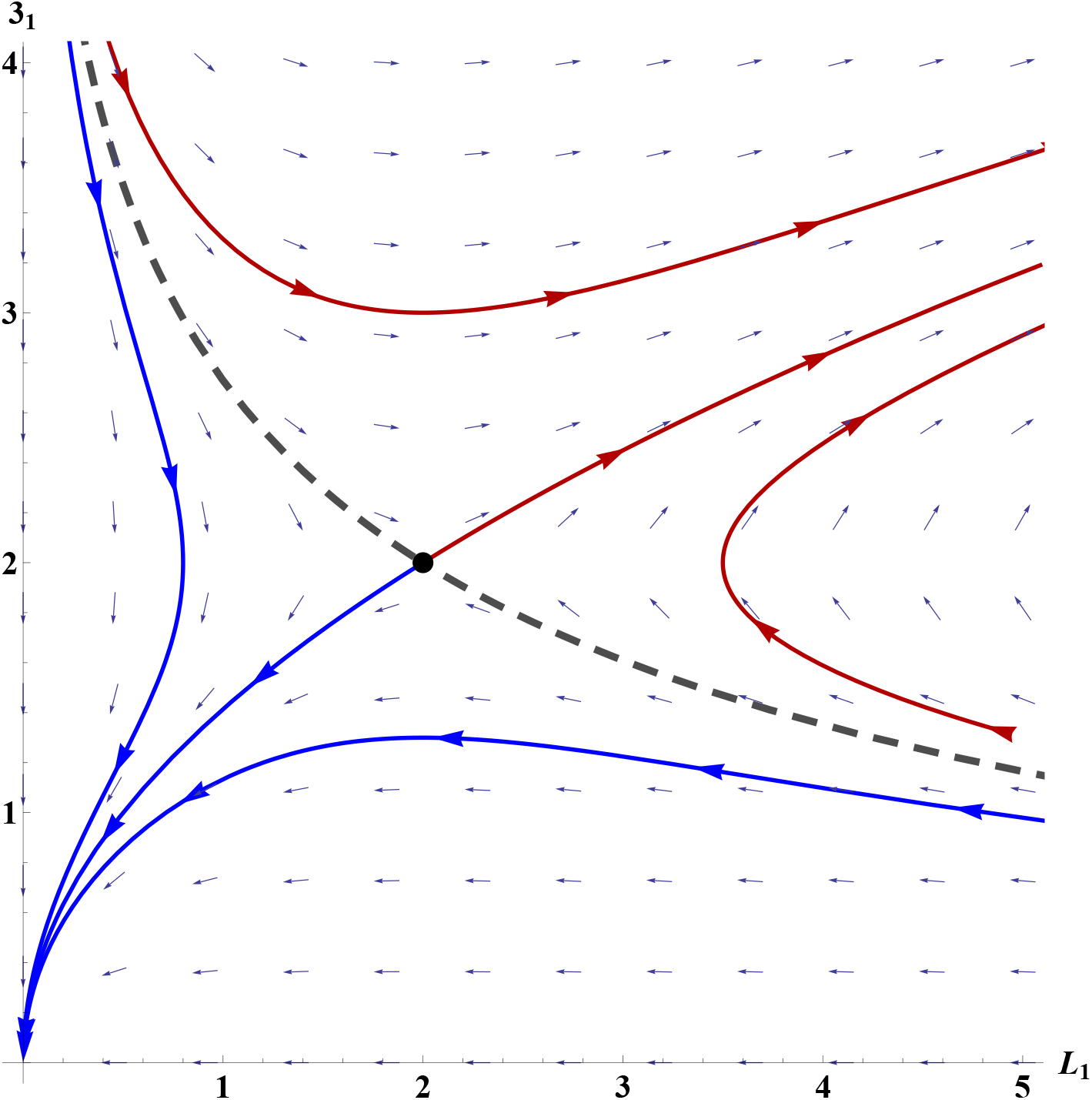
Phase plane for the toy model (Appendix B.1)–(Appendix B.2) for *σ* = *ρ* = 2. The qualitative behavior is similar to the four dimensional system (3.13)–(3.16). Note the two regions corresponding to trajectories converging to (0, 0) and trajectories diverging to infinity in both component. The two regions are separated by the stable manifold of the saddle point (dashed).

## 6 Discussion

In this paper, we have investigated the tissue phase separation event of the compaction step at the start of precartilage mesenchymal condensation in the chicken limb bud, along with the bistable dynamical switch that defines this transition. The switch constitutes the core of the reaction-diffusion-adhesion network that regulates pattern-formation of the limb skeleton in this species. The core module controls the transition between two lineage-adjacent, behaviorally different cell states: uncondensed and protocondensed (Bhat et al. (2011)) precartilage mesenchyme. The phase space of the dynamical system consists of two regions separated by the stable manifold of a saddle point, one in which trajectories converge to a stationary state of non-expression of the protocondensation determinant Gal-1A and the other in which trajectories representing unbounded production of this molecule. The two fates thus correspond biologically to cells within a protocondensation and cells outside this region, which are marked, respectively (among other differences) by very high and very low levels of Gal-1A.

In addition to determining the concentrations of Gal-1A and Gal-8, the defining dynamical system for the cell state transition (3.13–3.14) specifies the concentrations of the receptor-bound forms of each of the two galectins, as well as the concentrations of the receptors themselves, one shared by the two galectins and a Gal-8-specific one. But while this set of molecules are controlling variables for the transition, they do not exhaustively define the intra- and extra-protocondensed states, which include differences in levels or activities of BMP receptor 1b, the transcription factor Sox9 (Barna and Niswander (2007)) and TGF-*β* and responsive nuclear factors that in the following hours will lead to expression of the fibronectin gene as definitive mesenchymal condensations are consolidated (Leonard et al. (1991)).

While the cell-state switch functions as part of a reaction-diffusion-adhesion network in the embryonic limb bud and high-density mesenchymal cell cultures derived from it, extracting it mathematically from this context allows us to ascertain some of its important features. First, however, limitations of such an operation must be acknowledged. The isolated core switching network from the model of Glimm et al. (2014) predicts that the Gal-1A concentration grows without bounds or goes to zero in the intra- and extra-protocondensation domains. These non-biological outcomes are artifacts of the isolation of a portion of the full network, in simulations of which realistic values of the galectins are predicted (Glimm et al. (2014); Bhat et al. (2019)). If the assumption of linear production of galectins and counterreceptors is replaced by sigmoidal production terms, the region of unbounded trajectories instead becomes the basin of attraction for a second steady state, and thus the system then displays typical bistability attained via a saddle-node bifurcation (Strogatz (1994); Griffith (1971); Graham et al. (2010)) with a “high” and a “low” state, see section 4.3.

More usefully, we note that attractor states of a given dynamical system are not necessarily conserved when components of the system can diffuse (Krause et al. (2018)), or even when identical copies of the system interact with each other (Kaneko and Yomo (1999)). It was therefore unclear before the present study whether or not the foci of high Gal-1A accumulation surrounded by regions of low abundance of this protein generated by the full reaction-diffusion-adhesion system (Glimm et al. (2014)) were dependent on cell-cell adhesion or communication. Our conclusion from the analysis here that the switch is, in fact, cell-autonomous suggests that it could have been a primitive feature of skeletogenic mesenchymal cells not dependent on fine-tuned, tissue-wide interactions. This, in turn, corresponds to evolutionary scenarios in which variously distributed nodules and plates of endoskeletal tissue arose in the fins of ancestral chordates prior to the stereotypical patterning systems seen in the appendages of sarcopterygian fishes, and particularly tetrapods (Newman et al. (2018)).

Another significant finding is that Gal-1A production is not susceptible to entering an oscillatory state within the biologically plausible region of state space we have explored. This is consistent with the results of a previous study in which a periodic function representing the experimentally ascertained oscillatory behavior of the transcriptional regulator Hes1 was incorporated into the full reaction-diffusion-adhesion system where it was postulated to control the cell adhesive response to Gal-1A (Bhat et al., 2019). Notwithstanding the potential entrainment of Gal-1A synthesis by this effect, the accumulation of the galectins increases monotonically under a range of simulation conditions in the Hes1-enhanced model as it did over the first 36 h of development in vitro (Bhat et al. (2019)).

While Hes1 oscillations and their synchronization appear to play an important role in the spatial regularity and refinement of the boundaries between condensed and non-condensed mesenchyme in the developing limb (Bhat et al. 2019), the circuitry described here that robustly keeps the production of Gal-1A from becoming periodic may be advantageous in a developmental mechanism since the opposite would lead to discontinuities in the spatial pattern (see Glimm et al., 2014; Newman et al., 2021).

We characterized the relation between cell density and Gal-1A distribution to verify the autonomy of core switching circuit from a key property of the spatiotemporal pattern forming system from which it was mathematically extracted. In a previous study, we showed that a patterned distribution of Gal-1A does not form in the absence of the advection component, i.e., movement of cells up the Gal-1A adhesion gradient specified by the 2GL reaction-diffusion-adhesion network (Glimm et al. (2014)). That is, the pattern-forming mechanism is “morphodynamic” in the sense of requiring cell or tissue rearrangement concurrently with the chemical signaling that drives the rearrangement (Salazar-Ciudad et al. (2003)). This raised the possibility that the elevated Gal-1A in protocondensations in the living system was tied to the elevated cell density at these sites. However, our quantitative analysis shows that the inherent properties of the core circuit, which lacks not only cell movement but even a cell-cell adhesive function, ensures that Gal-1A concentration is higher on a per cell basis inside than outside protocondensations (Fig. 9).

Our observation of greater motility of mesenchymal cells inside forming limb condensations relative to those outside is at odds with the classical model of haptotaxis, in which increased cell density is a function of enhanced adhesivity (Dickinson, 2000), particularly when considering the higher levels of Gal-1A within protocondensations. In light of the changed motile behavior and reduced cell-substratum interaction of the condensing cells, and their confinement within a sharp perimeter, we propose that the tissue undergoes a transition to a more fluid state (Jain et al., 2020; Kim et al., 2021). Protocondensation formation can then be interpreted as the separation of a fluid from a semi-solid material.

Certain extracellular biomolecules such as Galectin-3 (a member of the same protein family as Gal-1A and Gal-8) induce liquid-like behaviors at cell surfaces due to intrinsic disorder and moderate-to-low ligand binding affinities (Zhao et al. (2021); Pally and Bhat (2021)). Given the flexibility of the Gal-8 linker domain (Earl et al. (2011); Si et al. (2016)) and the propensity for Gal-1 and Gal-8 to aggregate and mediate low avidity protein-glycan binding, it is reasonable to hypothesize that high expression of one or both of these galectins within forming condensations ‘trap’ cells without impeding their motility. Fibronectin, another ECM glycoprotein that accumulates during condensation (Frenz et al., 1989) may also promote the formation of this fluidized mesenchymal phase (Palamidessi et al. (2019); Gopal et al. (2017)).

The formation of limb bud protocondensations by focal tissue fluidization is a developmentally facile way of setting up a patterned array of nodules, but if a skeleton is to result, the elements must ultimately become solidified. In present-day species this occurs by conversion of the primordia to cartilage (later typically replaced by bone) in tetrapods, or directly to bone in the dermal rays of ray-finned fishes. This leads us to propose that formation of fluid domains might have been an evolutionarily ancestral mesenchymal patterning system (possibly not originally used for skeletogenesis) which subsequently recruited the transcription factors Sox9 or Runx2, respectively the cartilage and bone master regulators (reviewed in Newman et al., 2018). In keeping with our findings on the regulatory dynamics of fluidization, we suggest that additional evolutionary steps accompanying the formation of the tetrapod-type limb skeleton might have involved suppression of any propensity of the bistable switching mechanism to oscillate, so that the spatial control of the pattern by additional developmental signals could take place in tissue domains free from periodic perturbations.

## Acknowledgements

B.K. was partially supported by the National Science Centre grant 2016/21/B/ST1/03071. R.B. acknowledges support from DBT [BT/PR26526/GET/119/92/2017] and the Wellcome Trust/DBT India Alliance Fellowship/Grant [Grant Number IA/I/17/2/503312]. Prof. James Glazier participated in the analysis presented in Table 2.

## Appendix A: Methods

### Appendix A.1 Cell tracking experiment (section 2)

#### Cell Culture

We obtained fertile white Leghorn chicken eggs from Avian Services, Inc. (Frenchtown, NJ). We cut developing legs at Hamburger-Hamilton stage 25 (Hamburger and Hamilton, 1952) 0.3 mm from the distal end of the limb bud. We used the tips, the mesodermal component of which consists almost entirely of precartilage cells (Newman et al., 1981; Brand et al., 1985), to prepare cells for culture. Cultures were prepared essentially as described (Downie and Newman, 1994). Briefly, we dissociated the cells in 1% trypsin-EDTA (Sigma), filtered them through Nytex 20-*µ*m mono-filament nylon mesh (Tetko, Briarcliff Manor, NY), and resuspended them at 2.0 × 10^7^ cells/ml in defined medium (DM, Paulsen and Solursh, 1988) containing 10% fetal bovine serum. Filtration removed most of the limb-bud ectoderm, which remained in sheets after trypsinization. We deposited one to three cell spots (10 *µ*l each, containing 2.0 × 10^5^ cells) in a Falcon 60 mm tissue-culture dish (Falcon No. 351008) and allowed the cells to adhere to the dish at 38.5°C in a 5% CO_2_ atmosphere for 15 min before flooding the dish with serum-free DM. Although the initial cell plating density is above confluency, adding the medium disperses cells not attached to the substratum, resulting in a tightly-packed monolayer. We returned the cell cultures to the incubator for 24 hours, then sealed the lids of the culture dishes with Parafilm to prevent dehydration during time-lapse photomicrography.

#### Immunofluorescence

High density micromass cultures were fixed with ice-cold 100% methanol, washed with PBS and permeabilized with 0.02% Triton X-100 for 10 min. The cultures were incubated with affinity-purified polyclonal rabbit antiGal-1A antibody for 2 h at room temperature, followed by incubation with DyLight 594-conjugated secondary goat anti-rabbit antibody for 1h. Following counterstaining of DNA with DAPI, the cultures were imaged with an inverted Zeiss IM35 epifluorescence microscope with appropriate filters and a high-magnification objective (63 **r***A* ∼ oil immersion lens). Post image acquisition, the images were grayscaled and intensity measured using Matlab.

#### Microscopy and Image Acquisition

We placed the culture dish in a temperature-controlled Peltier warming device, maintained at 38.5deg C on the stage of a Zeiss IM35 inverted microscope equipped with a 32× Planacromat phase-contrast objective. Warm air blown across the dish lid prevented condensation on the lid’s inner surface. A timer box turned the microscope’s light source on and off in synchrony with the computer-controlled camera to minimize cells’ exposure to light. We took time-lapse images at intervals of 2 minutes at a resolution of 640× 480 pixels. Of our eight experimental plates, three developed condensations near the center of the microscope’s field of view and therefore permitted quantitative study. We chose one of these cultures for detailed analysis. The others developed at the same rate and had the same general morphology as the one we analyzed. We collected more than 350 individual images of the selected culture. Cells in frames 241-350 increasingly defocused as differentiation to cartilage changed cell shapes and the culture thickened. In these frames we could not track individual cells, especially in the condensation center, so we processed and analyzed only frames 1-240.

#### Image Processing and Data Analysis for cell-tracking experiments

The resolution of the images was 0.285*µ*m*/*pixel, sufficient to track the 10*µ*m diameter cells’ center-of-mass motions and deformations. With the aid of ImageJ software, which can play a sequence of images forward and backward, we hand-traced the boundaries between individual cells using Photoshop. Cells moved much less than one cell radius between frames, so we could unambiguously identify which cells in each image corresponded to the same experimental cell. We then used Photoshop and Matlab to assign a district gray level to each experimental cell for display and tracking (Fig.1).

For the statistical analysis, we digitized every tenth image, processing 25 images. In the image set the interval between two successive processed images is 20 min and the total duration is 480 min.

We labeled only cells lying completely within the field of view during the entire time series. We did not label or use cells which crossed the boundary of the field of view or which moved in or out of the field of view during the series. We observed 27 cell divisions during the 760 min time-lapse series in a population of about 145 cells; 8 of 92 labeled cells divided during the first 480 min. When a cell divided, we assigned to one of its daughter cells its mother cell’s gray level and used it in our statistical measurements. We assigned the other a new gray level and omitted it from our statistical measurements. By these criteria, the image set has 92 cells.

We used Matlab for image processing and data analysis. We recorded two-dimensional cell surface areas, which we interpreted as areas of contact with the underlying substratum, for all labeled cells. We used the coordinates of the center of mass 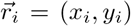 to analyze cell displacements. We recorded the mean-squared displacement, 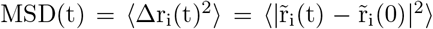, the average distance a cell’s center of mass traveled. We approximated the velocity of cells at time *t* as 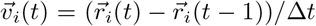, where Δ*t* is 20 min. We also measured the autocorrelation of velocities: 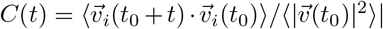, which quantifies the time scale over which cells maintained the magnitude and direction of their velocities.

## Appendix B: Illustrative simplified version of the system (3.17)– (3.19)

To further aid visualization of the three-dimensional system (3.17)–(3.19), illustrated in Figure 5, we can consider an even simpler two component system which nevertheless displays some of the same key behaviors as the three-component system. For this, note that for *π* = 1,

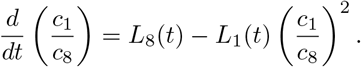

This is a nonautonomous equation for *u*(*t*) = *c* (*t*)*/c* (*t*) which then satisfies 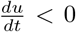 for 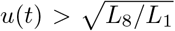 and 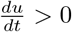 for 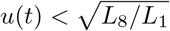. Motivated by this and the fact that *L*_8_(*t*) ⟶ 1*/σ* as *t* ⟶ ∞, we set up an illustrative toy model by taking the equations (3.13) and (3.15) for *L*_1_ and *c*_1_ and replace *c*_8_(*t*) by 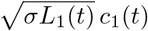. With this substitution, one obtains the two-component system

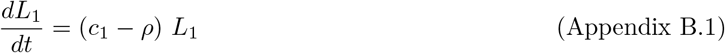

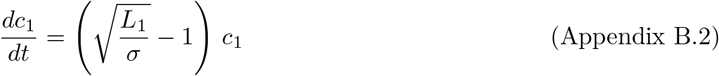

A typical phase plane is shown in Figure 10. The phase plane shows some of the same key behaviors as the one for the four component system (3.13)–(3.16), namely the separation of phase space into two regions corresponding to different cell fates.

Note that the system (Appendix B.1)–(Appendix B.2) is very similar to simple Lotka-Volterra type population models of mutualism with negative internal growth rates, representing populations that depend on each other for mutual survival (Murray (2002)).

## Appendix C: Proof of Proposition 4.5

We prove this result by matched asymptotic expansion. Let *r* = *πρ*; so we are interested in the case *r* → ∞. By (4.2), the pair of complex eigenvalues *λ*(*ρ, π*) and 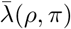 are solutions of the polynomial equation

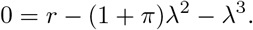

We seek an expansion in the form

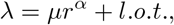

where “l.o.t.” denotes lower order terms in the power expansion in *r*. This results in the equation

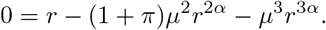

Comparing powers of *r*, this forces *α* = 1*/*3, and thus 0 = *µ*^3^ − 1, which gives the three root *µ*_1_ = 1, *µ*_2_ = exp(2*πi/*3), *µ*_2_ = exp(4*πi/*3). We thus obtain the expansion of the complex eigenvalues as

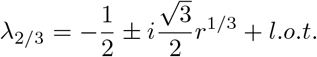

from where the desired result follows:

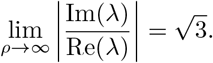

## Supplementary material

**Figure S1:**
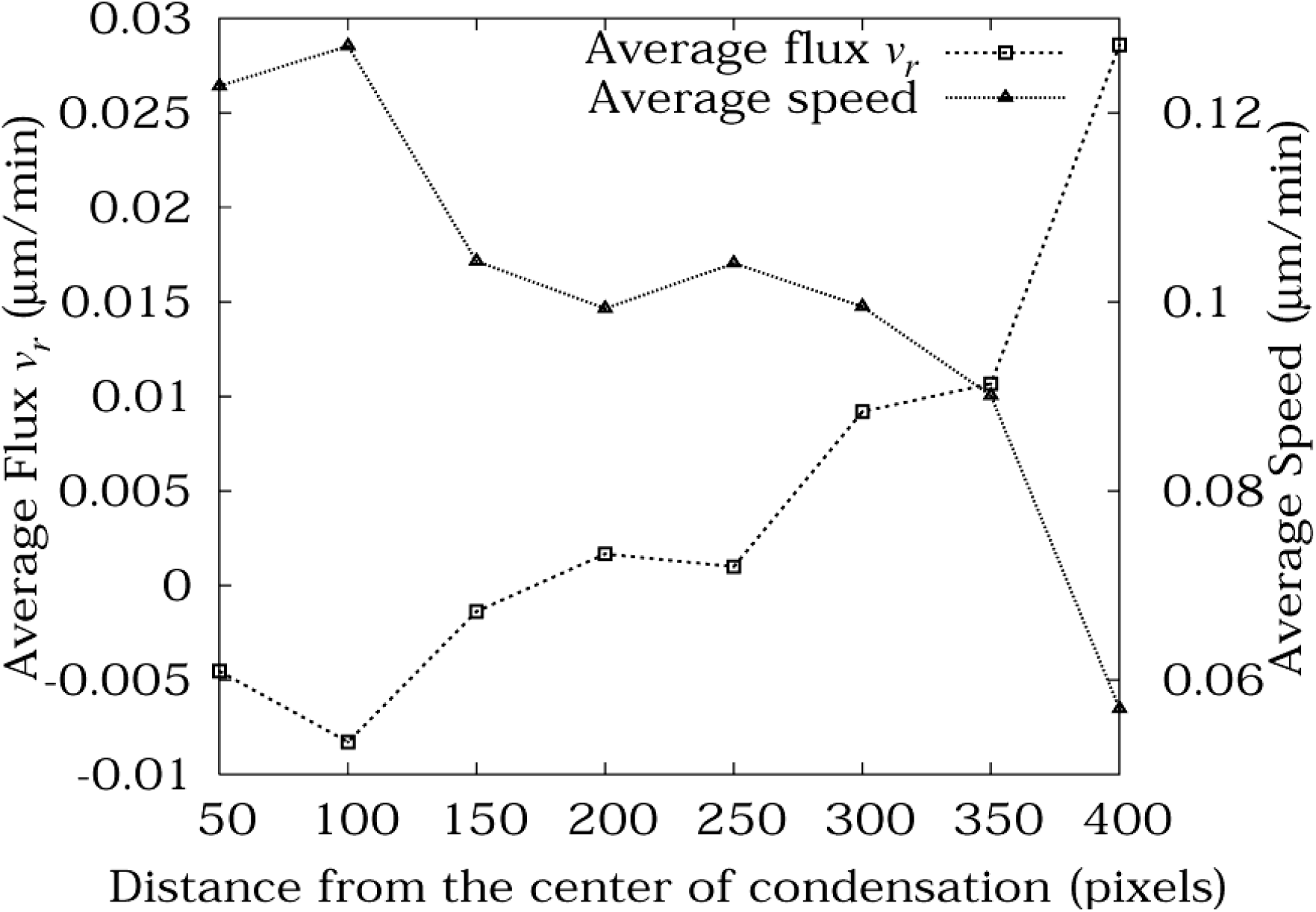
Cell flux towards the center of condensation and average cell speed as a function of distance from the center of condensation. Cell flux in a band is defined as *v*_*r*_ = **v** · **r** averaged over all cells in the band over all 25 images in the image set, where **v** is the velocity of the cell and **r** is the unit vector pointing from the center of mass of the cell to the center of condensation.

**Movie M1:** Time-lapse movie of the emergence of protecondenstion taken at 2 min intervals for 480 minutes total. (File cell•culture•seq•013103.mov)

## References

E. R. Alvarez-Buylla, A. Chaos, M. Aldana, M. Benítez, Y. Cortes-Poza, C. Espinosa-Soto, D. A. Hartasánchez, R. B. Lotto, D. Malkin, G. J. Escalera Santos, and P. Padilla-Longoria. Floral morphogenesis: stochastic explorations of a gene network epigenetic landscape. PloS one, 3(11):e3626, 2008. ISSN 1932-6203. doi: 10.1371/journal.pone.0003626.

M. Barna and L. Niswander. Visualization of Cartilage Formation: Insight into Cellular Properties of Skeletal Progenitors and Chondrodysplasia Syndromes. Developmental Cell, 12(6):931–941, June 2007. ISSN 1534-5807. doi: 10.1016/j.devcel.2007.04.016.

R. Bhat, K. M. Lerea, H. Peng, H. Kaltner, H. J. Gabius, and S. A. Newman. A regulatory network of two galectins mediates the earliest steps of avian limb skeletal morphogenesis. BMC Dev. Biol., 11:6, 2011.

R. Bhat, M. Chakraborty, T. Glimm, T. A. Stewart, and S. A. Newman. Deep phylogenomics of a tandem-repeat galectin regulating appendicular skeletal pattern formation. BMC Evolutionary Biology, 16(1):162, Aug. 2016. ISSN 1471-2148. doi: 10.1186/s12862-016-0729-6.

R. Bhat, T. Glimm, M. Linde-Medina, C. Cui, and S. A. Newman. Synchronization of Hes1 oscillations coordinates and refines condensation formation and patterning of the avian limb skeleton. Mechanisms of Development, 156:41–54, Apr. 2019. ISSN 0925-4773. doi: 10.1016/j.mod.2019.03.001.

H. H. Chang, P. Y. Oh, D. E. Ingber, and S. Huang. Multistable and multistep dynamics in neutrophil differentiation. BMC Cell Biology, 7(1):11, Feb. 2006. ISSN 1471-2121. doi: 10.1186/1471-2121-7-11.

C. Chen, C. A. Duckworth, B. Fu, D. M. Pritchard, J. M. Rhodes, and L.-G. Yu. Circulating galectins -2, -4 and -8 in cancer patients make important contributions to the increased circulation of several cytokines and chemokines that promote angiogenesis and metastasis. British Journal of Cancer, 110(3):741–752, Feb. 2014. ISSN 0007-0920. doi: 10.1038/bjc.2013.793.

F. Corson and E. D. Siggia. Gene-free methodology for cell fate dynamics during development. eLife, 6:e30743, Dec. 2017. ISSN 2050-084X. doi: 10.7554/eLife.30743. Publisher: eLife Sciences Publications, Ltd.

F. Corson, L. Couturier, H. Rouault, K. Mazouni, and F. c. Schweisguth. Self-organized Notch dynamics generate stereotyped sensory organ patterns in Drosophila. Science (New York, N.Y.), 356(6337), 2017. ISSN 1095-9203. doi: 10.1126/science.aai7407.

M. Cui, E. Vielmas, E. H. Davidson, and I. S. Peter. Sequential Response to Multiple Developmental Network Circuits Encoded in an Intronic cis-Regulatory Module of Sea Urchin hox11/13b. Cell Reports, 19(2):364–374, 2017. ISSN 2211-1247. doi: 10.1016/j.celrep.2017.03.039.

R. B. Dickinson. A generalized transport model for biased cell migration in an anisotropic environment. Journal of Mathematical Biology, 40(2):97–135, Feb. 2000. ISSN 1432-1416. doi: 10.1007/s002850050006.

L. A. Earl, S. Bi, and L. G. Baum. Galectin multimerization and lattice formation are regulated by linker region structure. Glycobiology, 21(1):6–12, Jan. 2011. doi: 10.1093/glycob/cwq144.

J. E. Ferrell. Bistability, bifurcations, and Waddington’s epigenetic landscape. Current biology: CB, 22(11): R458–466, June 2012. ISSN 1879-0445. doi: 10.1016/j.cub.2012.03.045.

D. A. Frenz, N. S. Jaikaria, and S. A. Newman. The mechanism of precartilage mesenchymal condensation: a major role for interaction of the cell surface with the amino-terminal heparin-binding domain of fibronectin. Developmental Biology, 136(1):97–103, Nov. 1989. ISSN 0012-1606. doi: 10.1016/0012-1606(89)90133-4.

C. Furusawa and K. Kaneko. A dynamical-systems view of stem cell biology. Science, 338(6104):215–217, Oct 2012.

T. Glimm, R. Bhat, and S. A. Newman. Modeling the morphodynamic galectin patterning network of the developing avian limb skeleton. J. Theor. Biol., 346:86–108, Apr 2014.

A. Goldbeter. Dissipative structures in biological systems: bistability, oscillations, spatial patterns and waves. Philosophical Transactions. Series A, Mathematical, Physical, and Engineering Sciences, 376(2124):20170376, July 2018. ISSN 1471-2962. doi: 10.1098/rsta.2017.0376.

D. K. Goode, N. Obier, M. S. Vijayabaskar, M. Lie-A-Ling, A. J. Lilly, R. Hannah, M. Lichtinger, K. Batta, M. Florkowska, R. Patel, M. Challinor, K. Wallace, J. Gilmour, S. A. Assi, P. Cauchy, M. Hoogenkamp, D. R. Westhead, G. Lacaud, V. Kouskoff, B. Gottgens, and C. Bonifer. Dynamic Gene Regulatory Networks Drive Hematopoietic Specification and Differentiation. Dev. Cell, 36(5):572–587, Mar 2016.

S. Gopal, L. Veracini, D. Grall, C. Butori, S. Schaub, S. Audebert, L. Camoin, E. Baudelet, A. Radwanska, S. Beghelli-de la Forest Divonne, S. M. Violette, P. H. Weinreb, S. Rekima, M. Ilie, A. Sudaka, P. Hofman, and E. Van Obberghen-Schilling. Fibronectin-guided migration of carcinoma collectives. Nature Communications, 8(1):14105, Jan. 2017. ISSN 2041-1723. doi: 10.1038/ncomms14105.

T. G. W. Graham, S. M. A. Tabei, A. R. Dinner, and I. Rebay. Modeling bistable cell-fate choices in the Drosophila eye: qualitative and quantitative perspectives. Development (Cambridge, England), 137(14): 2265–2278, July 2010. ISSN 0950-1991. doi: 10.1242/dev.044826.

J. S. Griffith. Mathematical neurobiology: an introduction to the mathematics of the nervous system. Academic Press, London, New York, 1971. ISBN 978-0-12-303050-4. Open Library ID: OL5758380M.

S. Huang. The molecular and mathematical basis of Waddington’s epigenetic landscape: a framework for post-Darwinian biology? Bioessays, 34(2):149–157, Feb 2012.

S. Huang, G. Eichler, Y. Bar-Yam, D. E. Ingber, and D. E. Ingber. Cell fates as high-dimensional attractor states of a complex gene regulatory network. Phys. Rev. Lett., 94(12):128701, Apr 2005.

A. Jain, V. Ulman, A. Mukherjee, M. Prakash, M. B. Cuenca, L. G. Pimpale, S. Münster, R. Haase, K. A. Panfilio, F. Jug, S. W. Grill, P. Tomancak, and A. Pavlopoulos. Regionalized tissue fluidization is required for epithelial gap closure during insect gastrulation. Nature Communications, 11(1), Nov. 2020. ISSN 2041-1723. doi: 10.1038/s41467-020-19356-x.

L. Jutras-Dubé, E. El-Sherif, and P. François. Geometric models for robust encoding of dynamical information into embryonic patterns. eLife, 9, Aug. 2020. ISSN 2050-084X. doi: 10.7554/eLife.55778.

K. Kaneko and T. Yomo. Isologous diversification for robust development of cell society. J. Theor. Biol., 199 (3):243–256, Aug 1999.

S. A. Kauffman. Metabolic stability and epigenesis in randomly constructed genetic nets. J. Theor. Biol., 22 (3):437–467, Mar 1969.

A. D. Keller. Specifying epigenetic states with autoregulatory transcription factors. Journal of Theoretical Biology, 170(2):175–181, Sept. 1994. ISSN 0022-5193. doi: 10.1006/jtbi.1994.1177.

S. Kim, M. Pochitaloff, G. A. Stooke-Vaughan, and O. Campàs. Embryonic tissues as active foams. Nature Physics, 17(7), July 2021. ISSN 1745-2481. doi: 10.1038/s41567-021-01215-1.

A. L. Krause, A. M. Burton, N. T. Fadai, and R. A. Van Gorder. Emergent structures in reaction-advection-diffusion systems on a sphere. Physical Review E, 97(4):042215, Apr. 2018. doi: 10.1103/PhysRevE.97.042215. Publisher: American Physical Society.

C. M. Leonard, H. M. Fuld, D. A. Frenz, S. A. Downie, J. Massague, and S. A. Newman. Role of transforming growth factor-β in chondrogenic pattern formation in the embryonic limb: Stimulation of mesenchymal con-densation and fibronectin gene expression by exogenenous TGF-β and evidence for endogenous TGF-β-like activity. Developmental Biology, 145(1):99–109, May 1991. ISSN 0012-1606. doi: 10.1016/0012-1606(91)90216-P.

M. Mojtahedi, A. Skupin, J. Zhou, I. G. Castaño, R. Y. Y. Leong-Quong, H. Chang, K. Trachana, A. Giuliani, and S. Huang. Cell Fate Decision as High-Dimensional Critical State Transition. PLOS Biology, 14(12): e2000640, Dec. 2016. ISSN 1545-7885. doi: 10.1371/journal.pbio.2000640. Publisher: Public Library of Science.

J. D. Murray. Mathematical biology. I, volume 17 of Interdisciplinary Applied Mathematics. Springer-Verlag, New York, third edition, 2002. ISBN 0-387-95223-3.

S. A. Newman. Cell differentiation: What have we learned in 50 years? Journal of Theoretical Biology, 485: 110031, Jan. 2020. ISSN 0022-5193. doi: 10.1016/j.jtbi.2019.110031.

S. A. Newman, T. Glimm, and R. Bhat. The vertebrate limb: An evolving complex of self-organizing systems. Progress in Biophysics and Molecular Biology, 137:12–24, Sept. 2018. ISSN 0079-6107. doi: 10.1016/j.pbiomolbio.2018.01.002.

S. A. Newman, R. Bhat, and T. Glimm. Spatial waves and temporal oscillations in vertebrate limb development. Biosystems, 208:104502, Oct. 2021. ISSN 0303-2647. doi: 10.1016/j.biosystems.2021.104502.

A. Palamidessi, C. Malinverno, E. Frittoli, S. Corallino, E. Barbieri, S. Sigismund, G. V. Beznoussenko, E. Martini, M. Garre, I. Ferrara, C. Tripodo, F. Ascione, E. A. Cavalcanti-Adam, Q. Li, P. P. Di Fiore, D. Parazzoli, F. Giavazzi, R. Cerbino, and G. Scita. Unjamming overcomes kinetic and proliferation arrest in terminally differentiated cells and promotes collective motility of carcinoma. Nature Materials, 18(11):1252–1263, Nov. 2019. doi: 10.1038/s41563-019-0425-1. Bandiera_abtest: a Cg type: Nature Research Journals Number:11 Primary_atype: Research Publisher: Nature Publishing Group Subject_term: Cell invasion;Tissues Subject_term_id: cell-invasion;tissues.

D. Pally and R. Bhat. N-terminal tail prolines of Gal-3 mediate its oligomerization/phase separation. Proceedings of the National Academy of Sciences of the United States of America, 118(25):e2107023118, June 2021. doi: 10.1073/pnas.2107023118.

M. Ross, G. Kaye, and W. Pawlina. Histology: A Text and Atlas with Cell and Molecular Biology. Lippincott Williams & Wilkins, 2002. ISBN 9780781751247.

I. Salazar-Ciudad, J. Jernvall, and S. A. Newman. Mechanisms of pattern formation in development and evolution. Development, 130(10):2027–2037, May 2003. ISSN 0950-1991, 1477-9129. doi: 10.1242/dev.00425. Publisher: The Company of Biologists Ltd Section: Review.

Y. Si, Y. Wang, J. Gao, C. Song, S. Feng, Y. Zhou, G. Tai, and J. Su. Crystallization of Galectin-8 Linker Reveals Intricate Relationship between the N-terminal Tail and the Linker. International Journal of Molecular Sciences, 17(12):2088, Dec. 2016. doi: 10.3390/ijms17122088.

D. Srivastava. Making or breaking the heart: from lineage determination to morphogenesis. Cell, 126(6): 1037–1048, Sep 2006.

A. Steinacher, D. G. Bates, O. E. Akman, and O. S. Soyer. Nonlinear Dynamics in Gene Regulation Promote Robustness and Evolvability of Gene Expression Levels. PLoS ONE, 11(4):e0153295, 2016.

S. Strogatz. Nonlinear dynamics and chaos: with applications to physics, biology, chemistry, and engineering. Studies in nonlinearity. Addison-Wesley Publ., Reading, Mass, 1994. ISBN 978-0-201-54344-5.

N. Suzuki, C. Furusawa, and K. Kaneko. Oscillatory protein expression dynamics endows stem cells with robust differentiation potential. PLoS ONE, 6(11):e27232, 2011.

M. Sáez, R. Blassberg, E. Camacho-Aguilar, E. D. Siggia, D. A. Rand, and J. Briscoe. Statistically derived geometrical landscapes capture principles of decision-making dynamics during cell fate transitions. Cell Systems, pages S2405–4712(21)00336–7, Sept. 2021. ISSN 2405-4720. doi: 10.1016/j.cels.2021.08.013.

R. Thom. Structural stability and morphogenesis. W. A. Benjamin, Inc., Reading, Mass.-London-Amsterdam, 1976. An outline of a general theory of models, Translated from the French by D. H. Fowler, With a foreword by C. H. Waddington, Second printing.

J. J. Tyson and B. Novak. Models in biology: lessons from modeling regulation of the eukaryotic cell cycle. BMC Biol., 13:46, Jul 2015.

J. J. Tyson, K. Chen, and B. Novak. Network dynamics and cell physiology. Nature Reviews Molecular Cell Biology, 2(12):908–916, Dec. 2001. ISSN 1471-0080. doi: 10.1038/35103078. Number: 12 Publisher: Nature Publishing Group.

J. J. Tyson, K. C. Chen, and B. Novak. Sniffers, buzzers, toggles and blinkers: dynamics of regulatory and signaling pathways in the cell. Current Opinion in Cell Biology, 15(2):221–231, Apr. 2003. ISSN 09550674. doi: 10.1016/S0955-0674(03)00017-6.

H. Xiao and R. Wu. Quantitative investigation of human cell surface N-glycoprotein dynamics †Electronic supplementary information (ESI) available: Two supplementary figures, and eleven supplementary tables. See DOI: 10.1039/c6sc01814a Click here for additional data file. Click here for additional data file. Chemical Science, 8(1):268–277, Jan. 2017. ISSN 2041-6520. doi: 10.1039/c6sc01814a. URL https://www.ncbi.nlm.nih.gov/pmc/articles/PMC5458730/.

Z. Zhao, X. Xu, H. Cheng, M. C. Miller, Z. He, H. Gu, Z. Zhang, A. Raz, K. H. Mayo, G. Tai, and Y. Zhou. Galectin-3 N-terminal tail prolines modulate cell activity and glycan-mediated oligomerization/phase separation. Proceedings of the National Academy of Sciences of the United States of America, 118(19):e2021074118, May 2021. doi: 10.1073/pnas.2021074118.

